# Polyunsaturated fatty acid analogues differentially affect cardiac Na_v_, Ca_v_, and K_v_ channels through unique mechanisms

**DOI:** 10.1101/772640

**Authors:** Briana M. Bohannon, Xiaoan Wu, Marta E. Perez, Sara I. Liin, H. Peter Larsson

## Abstract

The cardiac ventricular action potential depends on several voltage-gated ion channels, including Na_v_, Ca_v_, and K_v_ channels. Mutations in these channels can cause Long QT Syndrome (LQTS) which increases the risk for ventricular fibrillation and sudden cardiac death. Polyunsaturated fatty acids (PUFAs) have emerged as potential therapeutics for LQTS because they are modulators of voltage-gated ion channels. Here we demonstrate that PUFA analogues vary in their selectivity for human voltage-gated ion channels involved in the ventricular action potential. The effects of specific PUFA analogues range from selective for a specific ion channel to broadly modulating all three cardiac ion channels (N_aV_, C_aL_, and I_Ks_). In addition, PUFA analogues do not modulate these channels through a shared mechanism. Our data suggest that different PUFA analogues could be tailored towards specific forms of LQTS, which are caused by mutations in distinct cardiac ion channels, and thus restore a normal ventricular action potential.

## Introduction

The ventricular action potential is mediated by the coordinated activity of several different voltage-dependent ion channels (1). The rapid upstroke of the ventricular action potential is mediated by the activation of the voltage-gated Na^+^ channel, Nav1.5, which then rapidly inactivates. The activation of L-type voltage gated Ca^2+^ channels, Cav1.2, and influx of Ca^2+^ leads to a sustained depolarization, or plateau phase, and the contraction of the cardiac muscle. Inactivation of Cav1.2 channels along with the activation of the slow delayed-rectifier K^+^ channels, Kv11.1 (which generates the I_Kr_ current) and Kv7.1/KCNE1 (which generates the I_Ks_ current), work to promote repolarization of the cell membrane (2). Mutations of any of these ion channels (or channelopathies) could lead to Long QT Syndrome (LQTS), which is an arrhythmogenic disorder that predisposes the individual to potentially fatal cardiac arrhythmias (3, 4).

The Nav1.5 α-subunit contains four non-identical linked domains, DI-DIV. Each of these domains contain 6 transmembrane segments (S1-S6), where the S1-S4 segments make up the voltage-sensing domains (VSD) and S5-S6 segments make up the pore domains (PD). The S4 helix of each of the four domains contains a motif with positively charged amino acid residues which allow the S4 segment to detect and respond to changes in the membrane electric field, acting as the channel voltage sensor (5). The movement of these voltage sensors determines the voltage dependence of activation and inactivation, where activation of the DI-III S4s are suggested to promote activation and activation of DIV S4 segment is sufficient to induce voltage-dependent inactivation of Nav1.5 (6). The Nav1.5 α-subunit exists as a macromolecular complex with the accessory subunit β1. β1 is a single transmembrane spanning helix that modifies the kinetics of Nav1.5 channel activation and inactivation and can alter the pharmacology of the Nav1.5 α-subunit (7–9). Gain-of-function mutations of Nav1.5 increase Na^+^ currents and lead to LQTS Type 3 (LQT3) (10–12).

Like Nav1.5, the Cav1.2 α-subunit contains four linked domains, DI-DIV, where each domain consists of 6 transmembrane segments S1-S6. S1-S4 form the VSD, where S4 acts as the voltage sensor, and S5-S6 form the PD. Cav1.2 exists as a large macromolecular complex with the accessory subunits β3 and α2δ subunits that are important for membrane expression and alter channel activation and deactivation kinetics, respectively (13, 14). Cav1.2 undergoes both voltage-dependent inactivation and Calcium-dependent inactivation (15, 16) which allows it to regulate Ca^2+^ influx into the cardiomyocyte. Gain-of-function mutations of Cav1.2 increase Ca^2+^ currents and lead to Long QT Type 8 (LQT8) (17, 18).

The voltage-gated K^+^ channel, Kv7.1, along with the auxiliary subunit KCNE1, mediates an important repolarizing K^+^ current, I_Ks_ (19–21). The Kv7.1 α-subunit contains 6 transmembrane segments, S1-S6. S1-S4 comprise the VSD, where S4 contains several positively charged amino acid residues that allow S4 to act as the voltage sensor of Kv7.1. S5-S6 segments comprise the channel PD. Kv7.1 forms a tetrameric channel, where 4 Kv7.1 α-subunits arrange to form a functional channel. The auxiliary β-subunit KCNE1 drastically modulates Kv7.1 channel voltage dependence, activation kinetics, and single-channel conductance (22, 23). Loss-of-function mutations in the Kv7.1 α-subunit and KCNE1 *β*-subunit lead to reductions in I_Ks_ and can lead to LQTS Type 1 (LQT1) and Type 5 (LQT5) (24–28), respectively.

Polyunsaturated fatty acids (PUFAs) are amphipathic molecules that have been suggested to possess antiarrhythmic effects (29, 30). PUFAs are characterized by having a long hydrocarbon tail with two or more double bonds, as well as having a charged, hydrophilic head group (31). PUFAs, such as DHA and EPA, have been shown to prevent cardiac arrhythmias in animal models and cultured cardiomyocytes by inhibiting the activity of Na_v_ and Ca_v_ channels (30, 32–34). Specifically, DHA and EPA are thought to bind to discrete sites on the channel protein to stabilize the inactivated states of the Nav and Cav channels (32, 35). Since the voltage sensors of Nav and Cav channels are relatively homologous, it has been suggested that PUFAs act on the voltage-sensing S4 segments that control inactivation in these channels (30, 32). Our group has demonstrated that PUFAs and PUFA analogues also modulate the activity of the Kv7.1/KCNE1 channel and work to promote voltage-dependent activation of the I_Ks_ current (36–38). The mechanism through which PUFAs promote Kv7.1/KCNE1 activation is referred to as the lipoelectric hypothesis which involves the following: 1) the PUFA molecule integrates into the membrane via its hydrocarbon tail and 2) the negatively charged PUFA head group electrostatically attracts the positively charged S4 segment and facilitates the outward movement of S4, promoting I_Ks_ channel activation (36). Our group has also demonstrated that PUFAs and modified PUFAs exert a second effect on the pore of Kv7.1 through an additional electrostatic interaction with a lysine residue (K326) in the S6 segment (39). This electrostatic interaction between the negatively charged PUFA head group and K326 leads to an increase in maximal conductance of the channel (G_max_) (39).

Some groups have suggested that PUFAs could modify Na_v_ channels by causing a leftward shift in voltage dependent inactivation through an electrostatic effect on the voltage-sensing domains involved in inactivation (30, 32). It is possible that PUFAs modulate Kv7.1/KCNE1, Nav, and Cav channels by a similar mechanism by integrating next to the S4 voltage sensors and electrostatically attracting the voltage sensors toward their outward position. If PUFAs integrate preferentially next to the S4 that controls inactivation in Nav and Cav channels but next to all S4s in Kv7.1/KCNE1 channels, PUFAs would promote activation in Kv7.1/KCNE1 channels but promote inactivation in Nav and Cav channels. Though both PUFAs and PUFA analogues are known to modulate different ion channel activities (i.e. processes underlying activation and inactivation), it is unclear whether specific PUFAs and PUFA analogues are selective for certain ion channels or if they broadly influence the activity of several different ion channels simultaneously.

In this work, we characterize the channel-specific effects of different PUFAs and PUFA analogues in order to further understand which PUFAs and PUFA analogues would be the most therapeutically relevant in the treatment for different LQTS subtypes. We have found that PUFA analogues modulate Kv7.1/KCNE1, Cav1.2, and Nav1.5 through different mechanisms instead of through a shared mechanism. In addition, we demonstrate that PUFA analogues exhibit a broad range of differences in selectivity for Kv7.1/KCNE1, Cav1.2, and Nav1.5. Lastly, PUFA analogues that are more selective for Kv7.1/KCNE1 are able restore a prolonged ventricular action potential and prevent arrhythmia in simulated cardiomyocytes.

## Results

### PUFA analogues modulate Kv7.1/KCNE1, Cav1.2, and Nav1.5 through distinct mechanisms

There are several studies supporting electrostatic activation of Kv7.1/KCNE1 channels by PUFA analogues (37–39, 41). PUFAs are known to inhibit Na_v_ and Ca_v_ channels, but there is little evidence on the mechanism of channel inhibition using a diverse set of PUFA analogues. Previous groups have suggested that PUFAs may inhibit Na_v_ and Ca_v_ channels by interacting with S4 voltage sensors and stabilizing the inactivated state since there are similarities between the voltage sensor profiles of Na_v_ and Ca_v_ channels. (30, 32–34). For this reason, we hypothesize that PUFA analogues inhibit Cav1.2 and Nav1.5 through a shared electrostatic mechanism on S4 voltage sensors, similar to that reported with Kv7.1/KCNE1 channels. But in the case of Na_v_ and Ca_v_ channels, PUFAs would left-shift the voltage dependence of inactivation instead of activation which is seen in Kv7.1/KCNE1. To compare the effects of different PUFA analogues on these three different channels, we here measure the currents from Kv7.1/KCNE1, Cav1.2, and Nav1.5 expressed in *Xenopus* oocytes using two-electrode voltage clamp.

We first illustrate the effects of a representative PUFA analogue Linoleoyl taurine (Lin-taurine) on the voltage dependence of activation and the conductance of Kv7.1/KCNE1 (Fig. 1A-C). These effects are reflected in the tail current-voltage relationship where the effects on the voltage sensor are measured as a leftward shift in the voltage dependence of activation and the effects on the conductance are measured as a relative increase in the maximal conductance upon PUFA application (Fig. 1C). We also measure the effect of Lin-taurine on Cav1.2 and Nav1.5 channels (Figure 2-3). When we apply Lin-taurine to the Cav1.2 macromolecular complex (Fig. 2A), we see that Lin-taurine reduces Ca^2+^ currents in a dose-dependent manner (Fig. 2B-C). However, Lin-taurine reduces Ca^2+^ current without shifting the voltage-dependence of Cav1.2 activation (Fig. 2B-C; Supplemental Fig. 1). We use a depolarizing pre-pulse protocol to measure changes in voltage-dependent inactivation (Fig. 2D-E). When we measure the effects of PUFA analogues on voltage-dependent inactivation, we see again a decrease in Ca^2+^ currents, but surprisingly no shift in voltage-dependent inactivation (Fig. 2D-E). This suggests that PUFA analogues do not inhibit Cav1.2 channels through a shared electrostatic mechanism on S4 voltage sensors that shifts the voltage dependence of S4 movement, but rather through a mechanism that reduces either the number of conducting channels (potentially through an effect on the pore) or the maximum conductance of each channel. When we apply Lin-taurine to Nav1.5 (Figure 3A) and measure voltage-dependent activation, we see a dose-dependent inhibition of Na^+^ currents with no shift in the voltage dependence of activation (Fig. 3B-C; Supplemental Fig. 1). However, when we measured voltage-dependent inactivation of Nav1.5, we observed that PUFA analogues left-shift the voltage dependence of inactivation (Fig. 3D-E). In addition to the left-shift in voltage-dependent inactivation, we also observe a dose-dependent decrease in Nav1.5 currents (Fig. 3E). This suggests that PUFA analogues, while they do influence the voltage dependence of inactivation, may also have an additional effect on the conductance of Nav1.5, leading to the dose-dependent decrease in Na^+^ currents seen on top of the leftward shift of the voltage dependence of inactivation.

**Figure 1:**
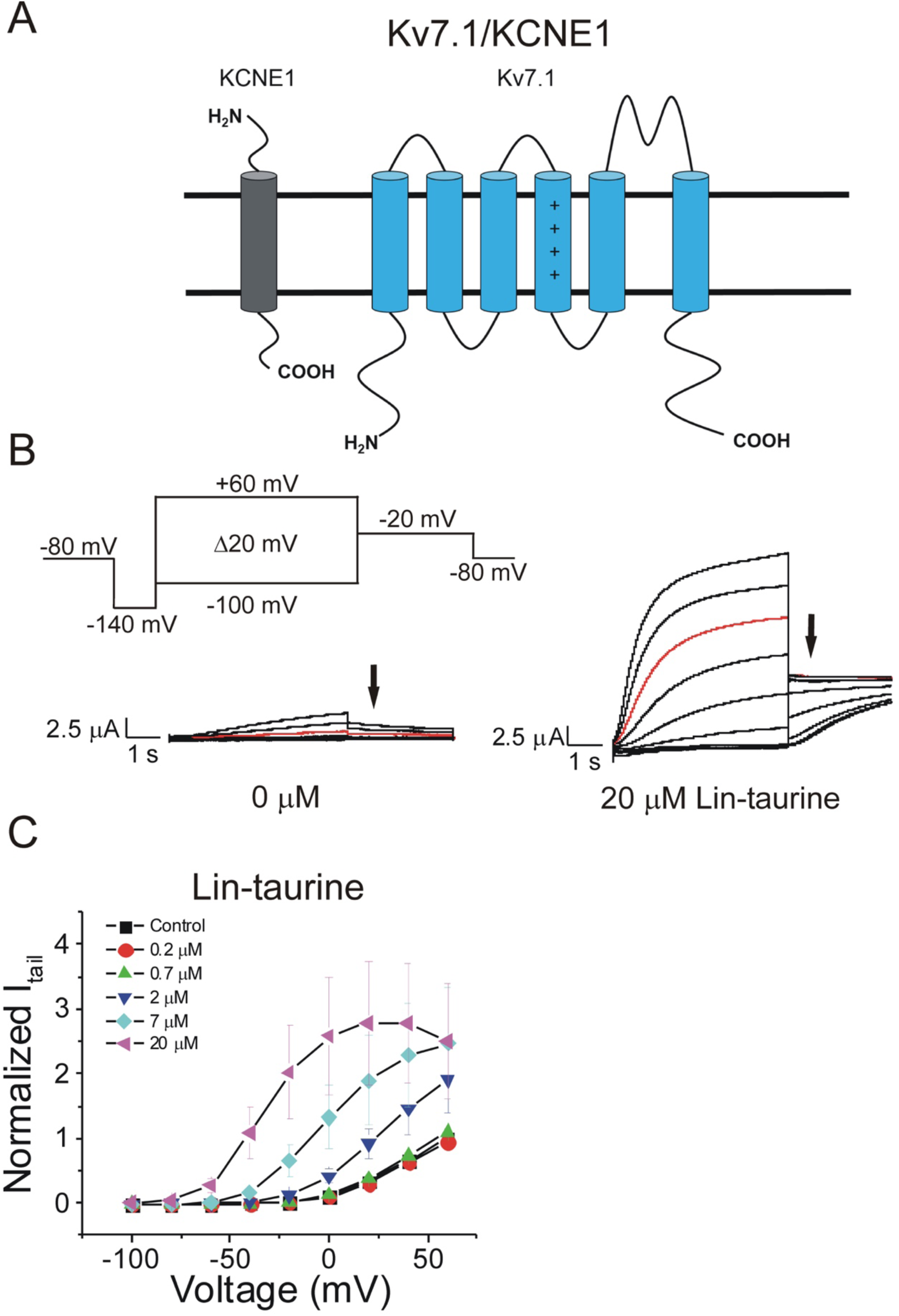
PUFAs activate Kv7.1/KCNE1 channels through an electrostatic mechanism on voltage sensor and pore. **A)** Simplified membrane topology of a single Kv7.1 α-subunit (blue) and a single KCNE1 β-subunit (grey). **B)** Voltage protocol used to measure voltage dependence of activation and representative Kv7.1/KCNE1 current traces in control (0 μM) and 20 μM Lin-taurine. Arrows mark tail currents. **C)** Current-voltage relationship demonstrating PUFA induced left-shift in the voltage-dependence of activation (V_0.5_) and increase in maximal conductance (G_max_) (mean ± SEM; n = 3).

**Figure 2:**
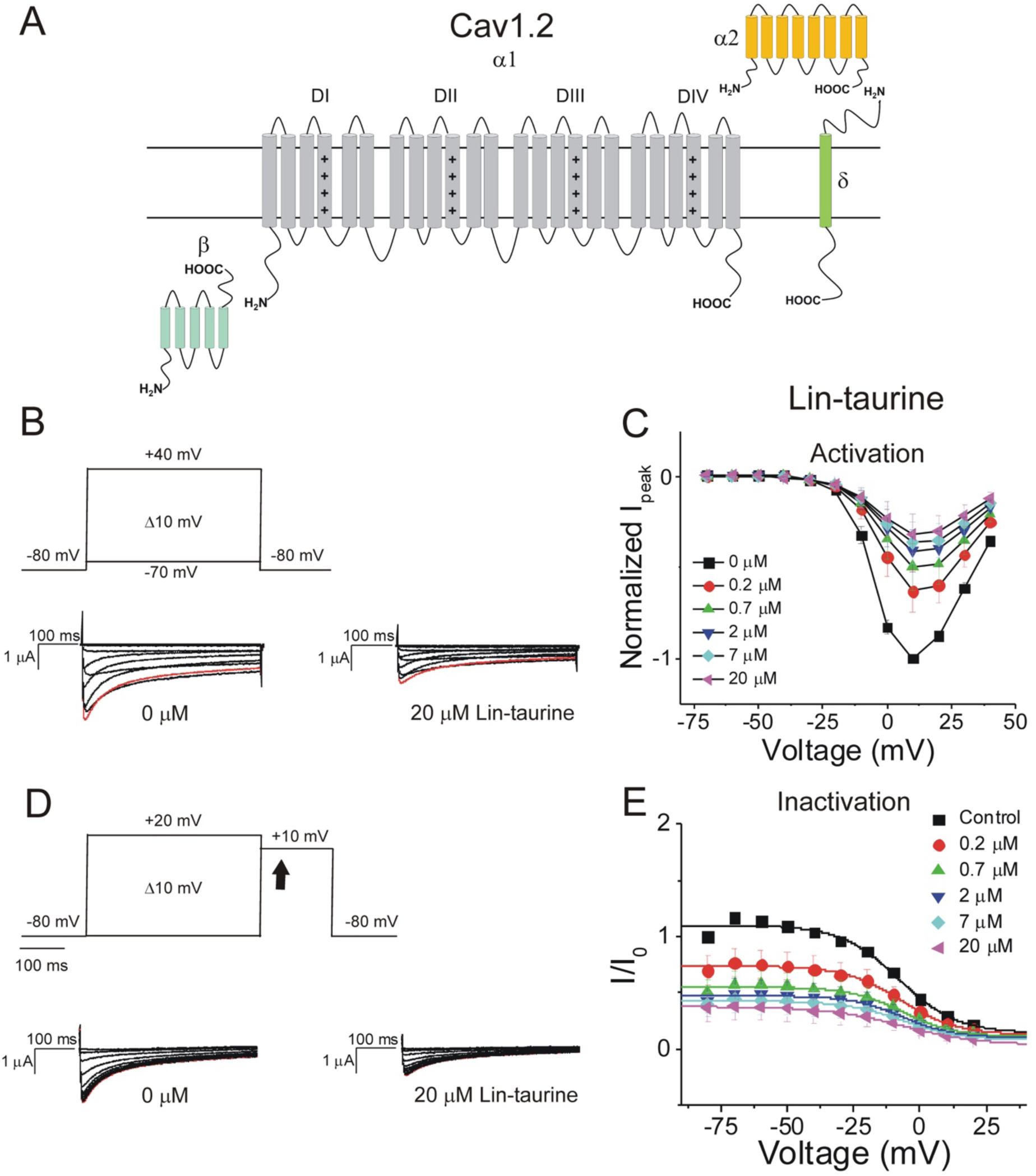
PUFAs inhibit Cav1.2 channels without altering channel voltage dependence. **A)** Simplified membrane topology of the Cav1.2 pore-forming α-subunit (light gray) and auxiliary β-(mint) and α2δ-subunits (yellow and green). **B)** Voltage protocol used to measure voltage dependence of activation and representative Cav1.2 current traces in control (0 μM) and 20 μM Lin-taurine. **C)** Current-voltage relationship demonstrating dose-dependent inhibition of Cav1.2 currents measured from activation protocol (mean ± SEM; n = 3). **D)** Voltage protocol used to measure voltage dependence of inactivation and representative Cav1.2 current traces in control (0 μM) and 20 μM Lin-taurine measured at arrow. **E)** Current-voltage relationship demonstrating dose-dependent inhibition of Cav1.2 currents measured from inactivation protocol (mean ± SEM; n = 3).

**Figure 3:**
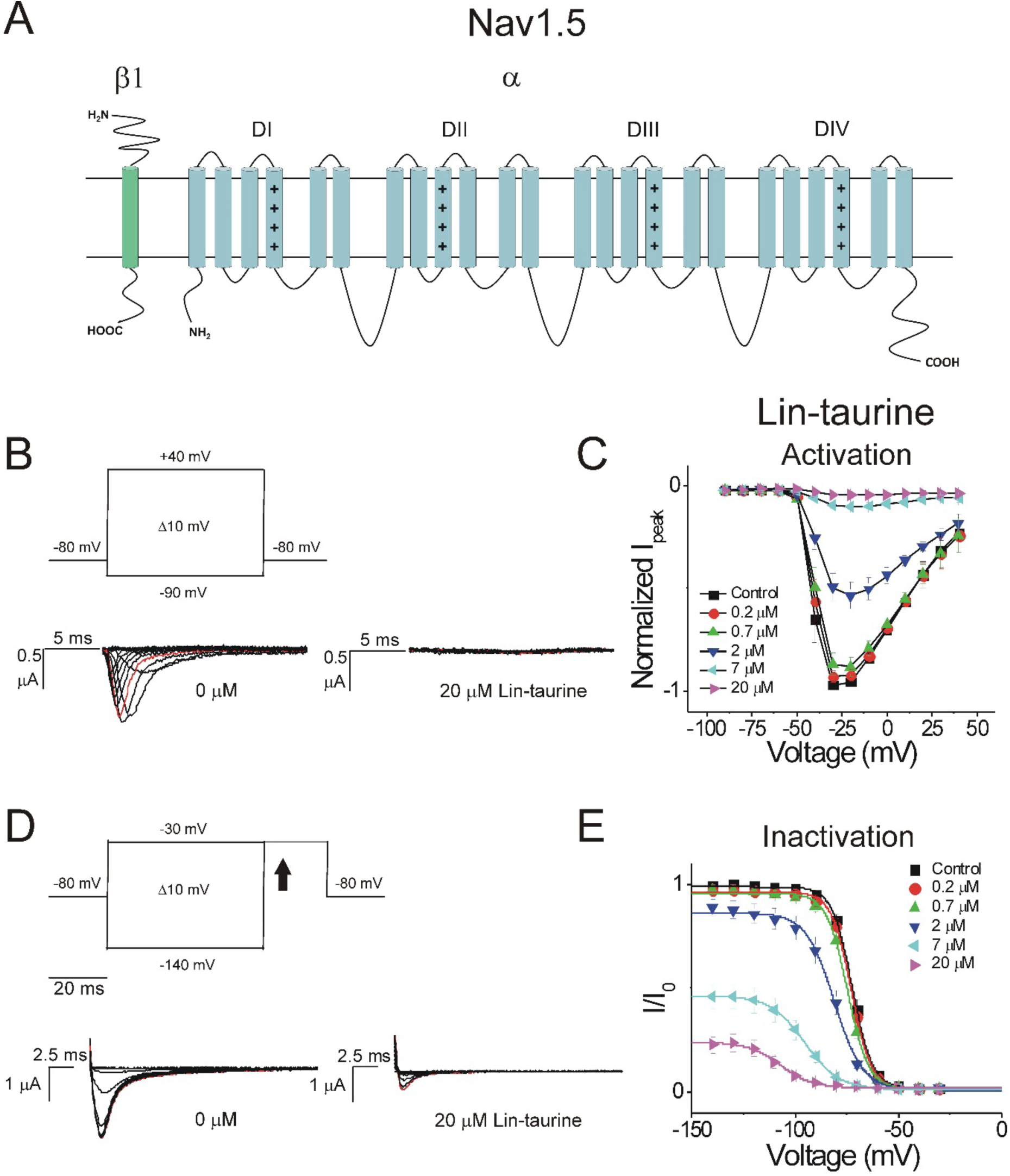
PUFAs inhibit Nav1.5 by shifting the voltage dependence of inactivation. **A)** Simplified membrane topology of the Nav1.5 pore-forming α-subunit (light blue) and auxiliary β-subunit (green). **B)** Voltage protocol used to measure voltage dependence of activation and representative Nav1.5 current traces in control (0 μM) and 20 μM Lin-taurine. **C)** Current-voltage relationship demonstrating dose-dependent inhibition of Nav1.5 currents measured from activation protocol (mean ± SEM; n = 5). **D)** Voltage protocol used to measure voltage dependence of inactivation and representative Nav1.5 current traces in control (0 μM) and 20 μM Lin-taurine measured at arrow. **E)** Current-voltage relationship demonstrating dose-dependent inhibition of Nav1.5 currents and leftward shift in the voltage dependence of inactivation measured from inactivation protocol (mean ± SEM; n = 5).

Through these data, we observe that PUFA analogues modulate cardiac voltage-gated ion channels through non-identical mechanisms. PUFA analogues promote the activation of Kv7.1/KCNE1 through electrostatic effects that left-shift the voltage dependence of activation and an increase in the maximal conductance. PUFA analogues inhibit Cav1.2 channels through an apparent effect on the pore leading to a reduction in Ca^2+^ current but without producing any leftward shift in the voltage dependence of inactivation. In addition, PUFA analogues inhibit Nav1.5 through a combination of a leftward shift in the voltage dependence of inactivation and an effect on the maximum conductance, which leads to a dose-dependent decrease in Na^+^ current. Together these findings show that PUFA analogues affect Kv7.1/KCNE1, Nav1.5, and Cav1.2 channels through different mechanisms.

### PUFA analogues with taurine head groups are non-selective and broadly modulate multiple cardiac ion channels, with preference for Cav1.2 and Nav1.5

We have found through previous work that PUFA analogues with taurine head groups are good activators of the Kv7.1/KCNE1 channels due to the low pKa of the taurine head group (37, 38). Having a lower pKa allows the taurine head group to be fully negatively charged at physiological pH so that it has maximal electrostatic effects on Kv7.1/KCNE1 channels (38). We tested a set of PUFA analogues with taurine head groups on Kv7.1/KCNE1, Cav1.2, and Nav1.5 channels to determine if these effects are selective for the Kv7.1/KCNE1 channel or if taurine analogues also modulate Cav1.2 and Nav1.5 channels. Lin-taurine is a PUFA analogue with a taurine head group (Figure 4A) that promotes the activation of the cardiac Kv7.1/KCNE1 channel, by promoting a leftward shift in the voltage-dependence of activation by −39.9 ± 3.6 mV at 7 μM (p = 0.008) (Fig 4B). In addition, the application of Lin-taurine produces a slight, but not statistically significant increase in the maximal conductance of the Kv7.1/KCNE1 channel at 7 μM (1.9 ± 0.6; p = 0.26) (Fig. 4C). Lin-taurine inhibits Cav1.2 current in a dose-dependent manner without left-shifting the voltage dependence of inactivation for Cav1.2 (2.8 ± 1.4 mV; p = 0.17), but instead by significantly decreasing the G_max_ at 7 μM (0.4 ± 0.1; p = 0.03) (Fig. 4D-E). Lastly, Lin-taurine inhibits Nav1.5 current by left-shifting the voltage dependence of inactivation (−23.5 ± 1.9 mV; p = 0.001) and also decreasing the G_max_ at 7 μM (0.5 ± 0.07; p = 0.005) (Fig 4F-G).

**Figure 4:**
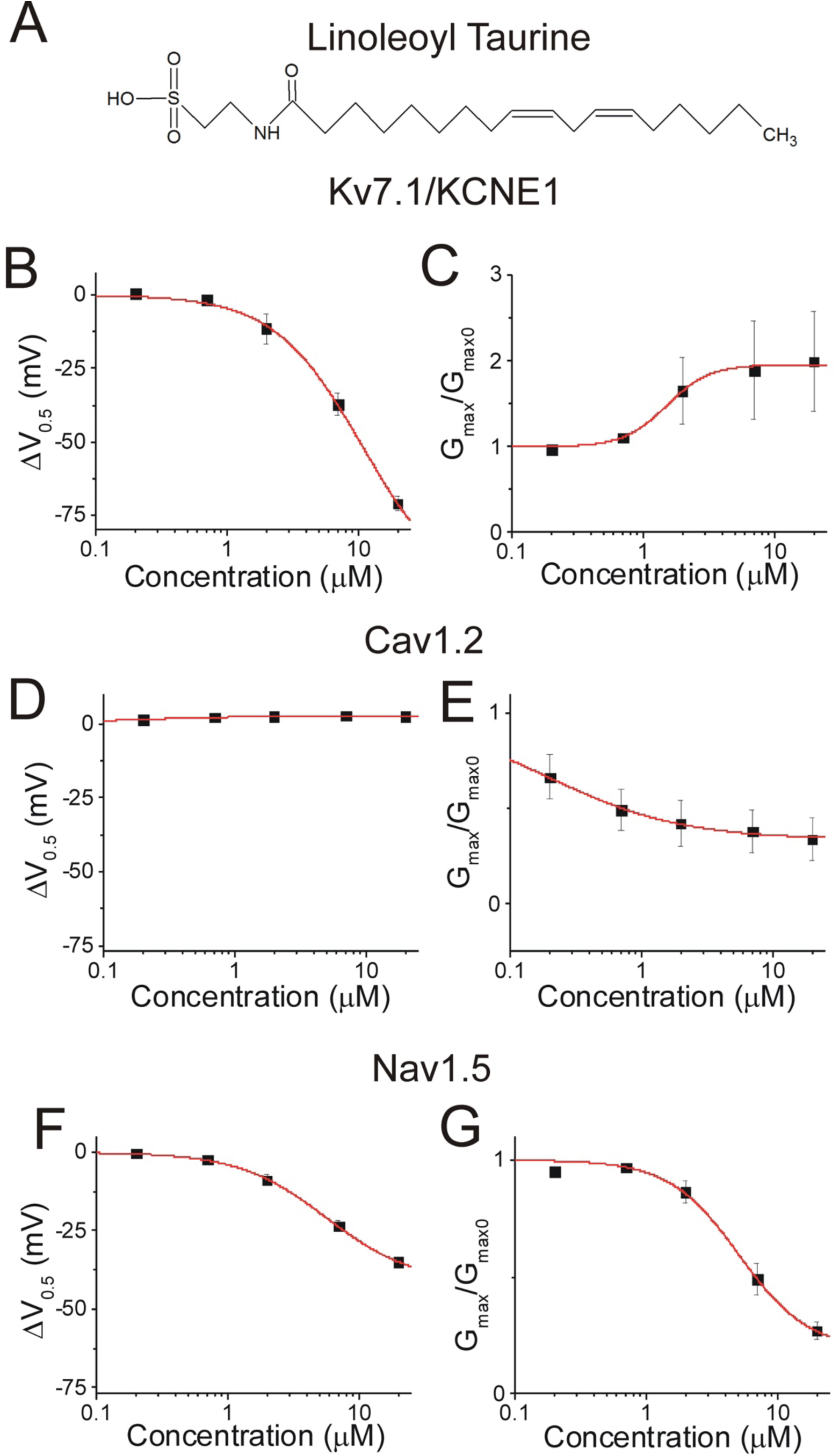
Linoleoyl taurine has broad selectivity for Kv7.1/KCNE1, Cav1.2, and Nav1.5. **A)** Structure of Linoleoyl taurine (Lin-taurine). **B, D, F)** Dose response of the shift in voltage dependent **B)** activation (ΔV_0.5_) of Kv7.1/KCNE1 channels (mean ± SEM; n = 3), **D)** inactivation (ΔV_0.5_) of Cav1.2 channels (mean ± SEM; n = 3), **F)** inactivation (ΔV_0.5_) of Nav1.5 channels (mean ± SEM; n = 5) in the presence of lin-taurine. **C, E, G)** Dose response of the change in maximal conductance (G_max_) of **C)** Kv7.1/KCNE1 channels **E)** Cav1.2 channels, **G)** Nav1.5 channels in the presence of lin-taurine.

N-arachidonoyl taurine (N-AT) is a PUFA analogue with a taurine head group that has been demonstrated by our group to promote activation of Kv7.1/KCNE1, left-shifting the voltage-dependence and increasing the G_max_ at 70 μM (Fig. 5A) (38). Here, we used lower concentrations (0.2, 0.7, 2, 7, and 20 μM) with the goal of understanding the selectivity of N-AT for cardiac ion channels and at more therapeutically feasible concentrations. Application of N-AT does not promote activation of Kv7.1/KCNE1 in this lower concentration range, does not left-shift of the voltage-dependence of activation (−1.8 ± 2.6 mV; p = 0.5), and does not increase the G_max_ at 7 μM (0.9 ± 0.03; p = 0.98) (Fig. 5B-D). However, N-AT causes a dose-dependent decrease in Cav1.2 current, though does not cause a significant shift in the voltage dependence of inactivation (13.5 ± 3.8 mV; p = 0.07), nor does it cause a significant reduction in the overall G_max_ (0.6 ± 0.1; p = 0.06) at 7 μM (Fig. 5E-G). In addition, N-AT decreases Nav1.5 current, produces a leftward shift in voltage-dependent inactivation (−16.7 ± 3.5 mV; p = 0.04), and significantly reduces the G_max_ at 7 μM (0.3 ± 0.04; p = 0.004) (Fig. 5H-J). This data suggests that N-AT is more selective for Cav1.2 and Nav1.5, compared to Kv7.1/KCNE1.

**Figure 5:**
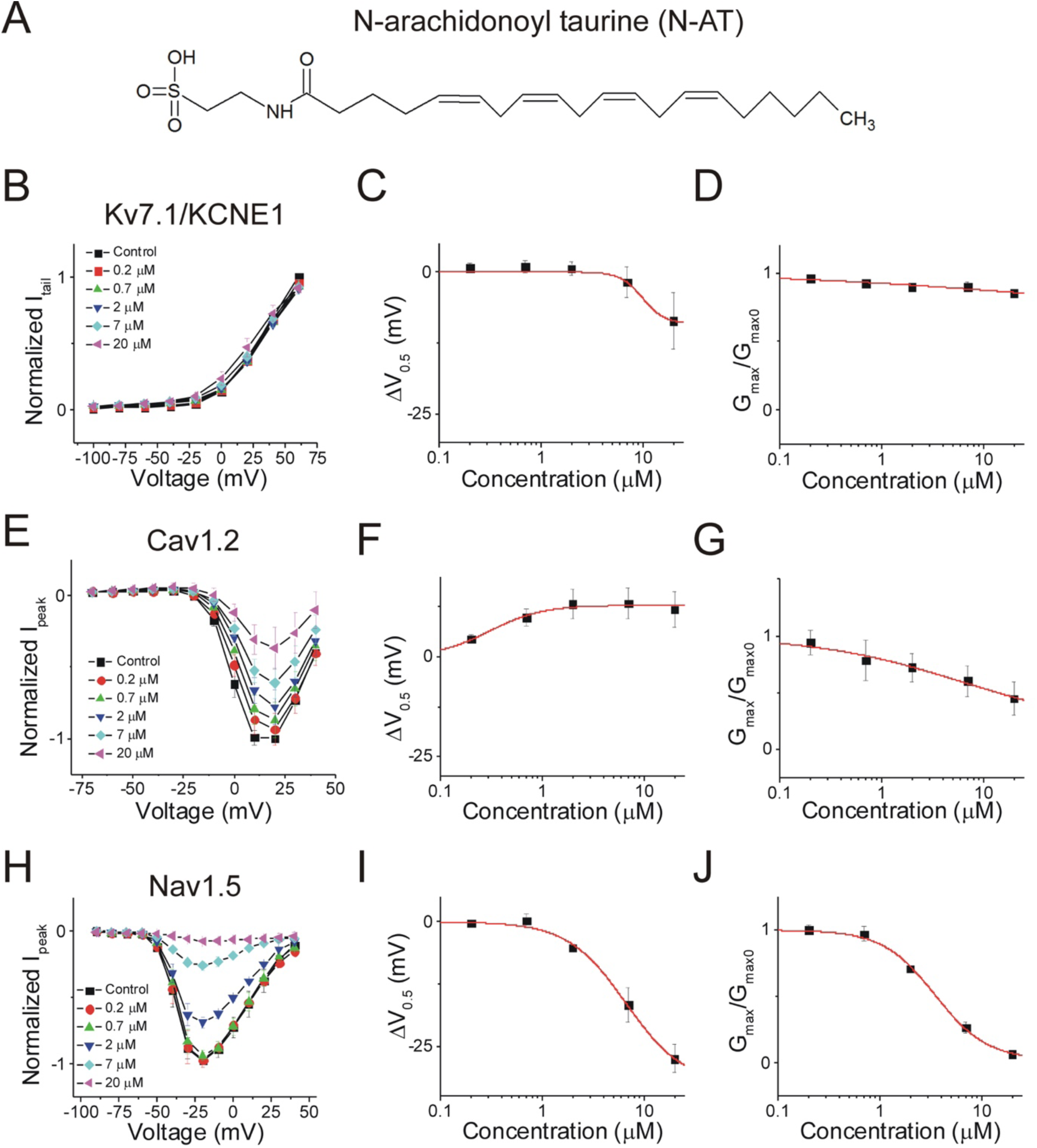
N-arachidonoyl taurine is more selective for Cav1.2 and Nav1.5 than for Kv7.1/KCNE1. **A)** Structure of N-arachidonoyl taurine (N-AT). **B)** Current-voltage relationship of N-AT on Kv7.1/KCNE1 channels (mean ± SEM; n = 5). **C)** Dose response of the shift in voltage dependent activation (ΔV_0.5_) of Kv7.1/KCNE1 channels in the presence of N-AT. **D)** Dose response of the change in maximal conductance (G_max_) of Kv7.1/KCNE1 channels in the presence of N-AT. **E)** Current-voltage relationship of N-AT on Cav1.2 channels (mean ± SEM; n = 4). **F)** Dose response of the shift in voltage dependent inactivation (ΔV_0.5_) of Cav1.2 channels in the presence of N-AT. **G)** Dose response of the change in maximal conductance (G_max_) of Cav1.2 channels in the presence of N-AT. **H)** Current-voltage relationship of N-AT on Nav1.5 channels (mean ± SEM; n = 3). **I)** Dose response of the shift in voltage dependent inactivation (ΔV_0.5_) of Nav1.5 channels in the presence of N-AT. **J)** Dose response of the change in maximal conductance (G_max_) of Nav1.5 channels in the presence of N-AT.

Pinoleoyl taurine (Pin-taurine) promotes the activation of Kv7.1/KCNE1 in a dose-dependent manner (Fig. 6A-B). Pin-taurine, like Lin-taurine, promotes a leftward shift in the voltage dependence of activation (−23.8 ± 2.7 mV; p = 0.003) and increases the G-max of Kv7.1/KCNE1 at 7 μM with a trend towards significance (2.2 ± 0.4; p = 0.06) (Fig. 6C-D). Pin-taurine also inhibits Cav1.2 current, but does not significantly shift the voltage-dependence of inactivation (4.1 ± 2.2 mV; p = 0.13) and does not significantly decrease the G_max_ at 7 μM (0.8 ± 0.1; p = 0.2) (Fig. 6E-G). Pin-taurine inhibits Nav1.5 currents and does so by significantly left-shifting the voltage dependence of inactivation (−16 ± 2.7 mV; p = 0.01) and decreasing the G_max_ at 7 μM (0.4 ± 0.09; p = 0.005) (Fig. 6H-J).

**Figure 6:**
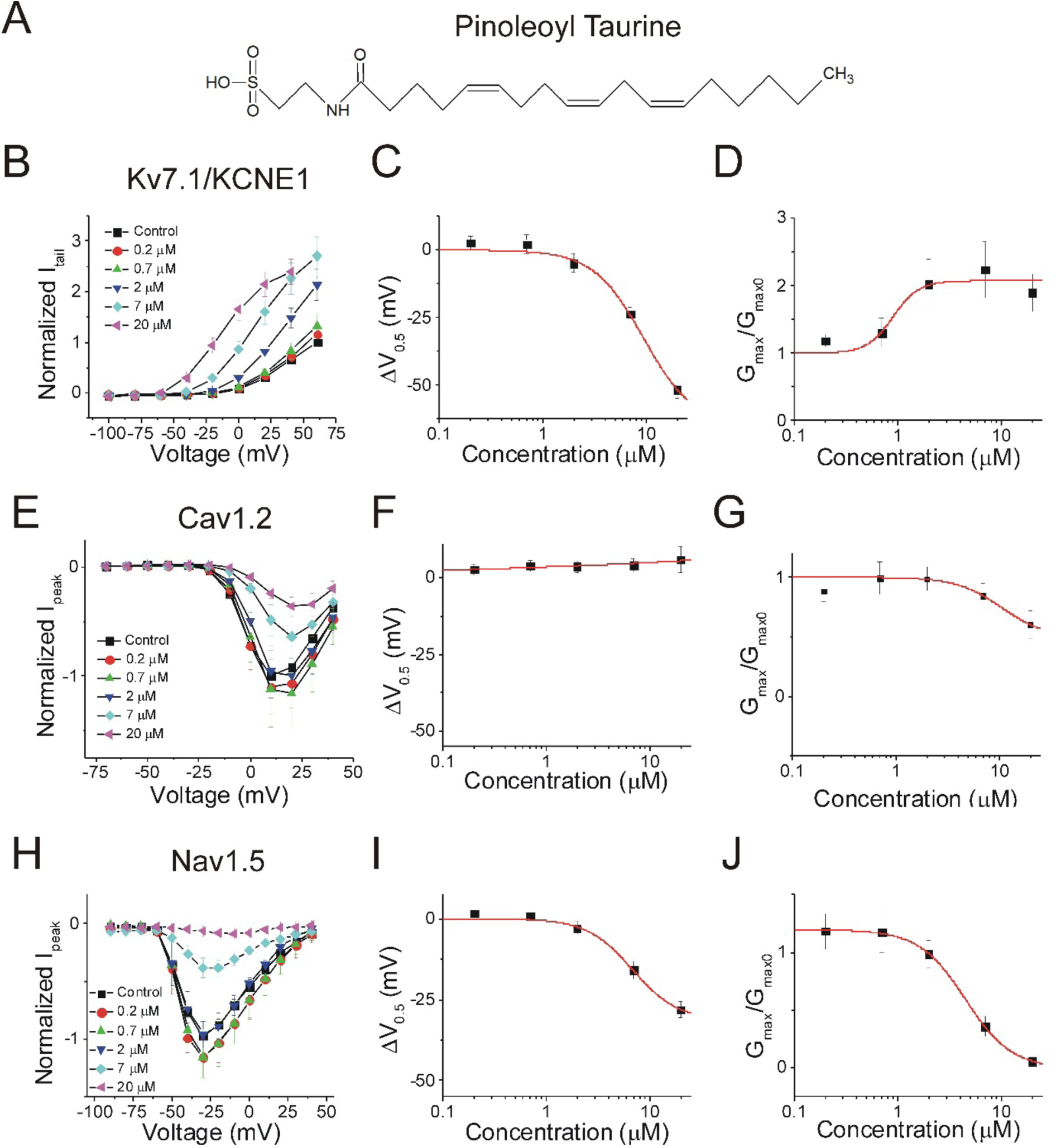
Pinoleoyl taurine has broad selectivity for Kv7.1/KCNE1, Cav1.2, and Nav1.5. **A)** Structure of Pinoleoyl taurine (Pin-taurine). **B)** Current-voltage relationship of pin-taurine on Kv7.1/KCNE1 channels (mean ± SEM; n = 4). **C)** Dose response of the shift in voltage dependent activation (ΔV_0.5_) of Kv7.1/KCNE1 channels in the presence of pin-taurine. **D)** Dose response of the change in maximal conductance (G_max_) of Kv7.1/KCNE1 channels in the presence of pin-taurine. **E)** Current-voltage relationship of pin-taurine on Cav1.2 channels (mean ± SEM; n = 5). **F)** Dose response of the shift in voltage dependent inactivation (ΔV_0.5_) of Cav1.2 channels in the presence of pin-taurine. **G)** Dose response of the change in maximal conductance (G_max_) of Cav1.2 channels in the presence of pin-taurine. **H)** Current-voltage relationship of pin-taurine on Nav1.5 channels (mean ± SEM; n = 4). **I)** Dose response of the shift in voltage dependent inactivation (ΔV_0.5_) of Nav1.5 channels in the presence of pin-taurine. **J)** Dose response of the change in maximal conductance (G_max_) of Nav1.5 channels in the presence of pin-taurine.

DHA-taurine promotes the activation of Kv7.1/KCNE1 channels in a dose-dependent manner, left-shifting the voltage-dependence of activation (−45.3 ± 2.9 mV; p = 0.004) and significantly increasing the G_max_ at 7 μM (1.7 ± 0.1; p = 0.03) (Fig. 7A-C). DHA-taurine application results in dose-dependent inhibition of Cav1.2 current (Fig. 7E), but does not significantly left-shift the voltage-dependence of inactivation at 7 μM (0.2 ± 0.8 mV; p = 0.85) (Fig. 7F). Instead, DHA-taurine causes a significant decrease in the G_max_ of Cav1.2 at 7 μM (0.4 ± 0.01; p < 0.001) (Fig. 7G). Lastly, DHA-taurine inhibits Nav1.5 by inducing a significant left-shift in the voltage-dependence of inactivation (−28.5 ± 0.6 mV; p < 0.001) and decreasing the G_max_ at 7 μM (0.05 ± 0.01; p < 0.001) (Fig. 7H-I).

**Figure 7:**
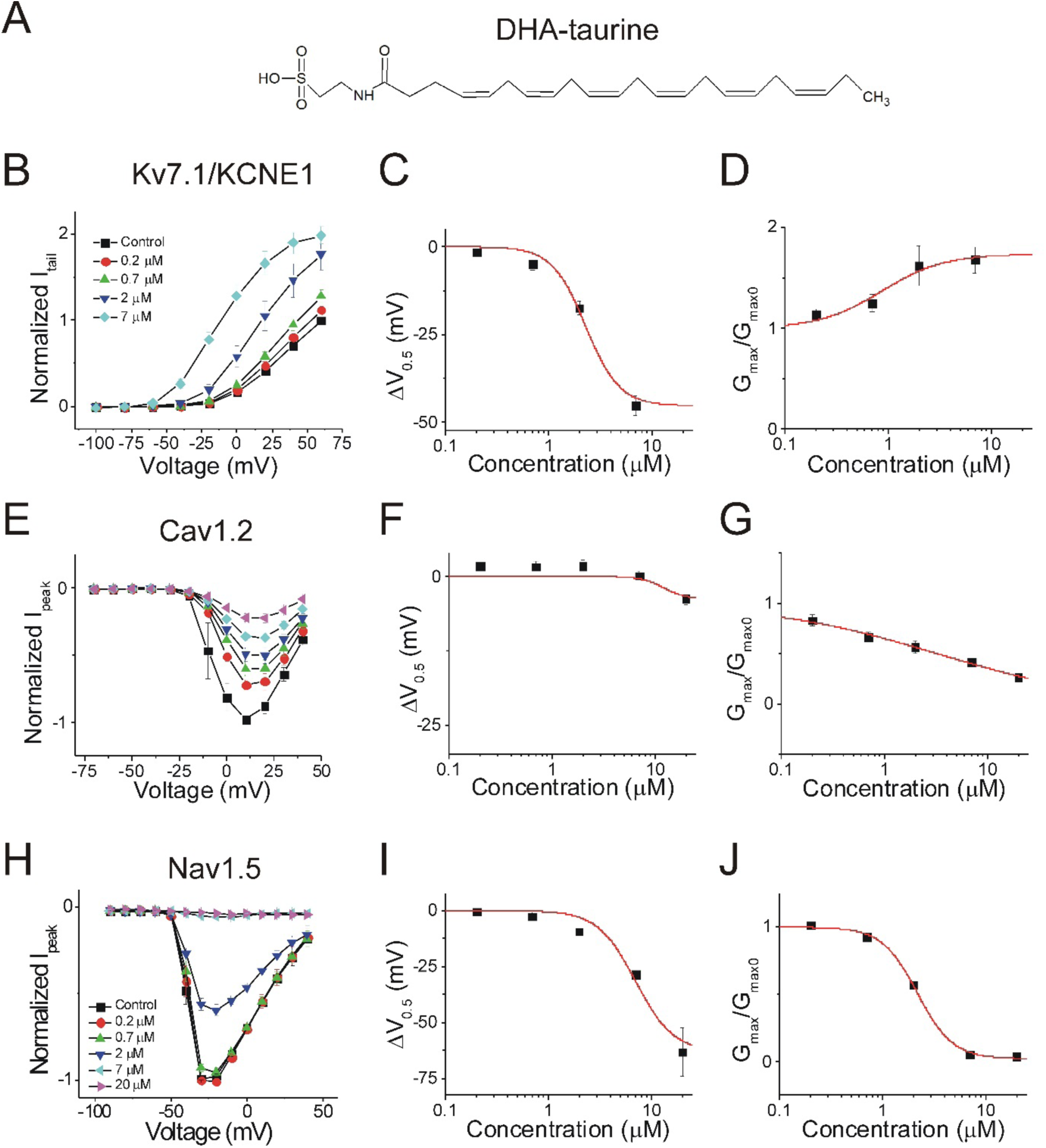
Docosahexanoyl-taurine has broad selectivity for Kv7.1/KCNE1, Cav1.2, and Nav1.5. **A)** Structure of docosahexanoyl taurine (DHA-taurine). **B)** Current-voltage relationship of DHA-taurine on Kv7.1/KCNE1 channels (mean ± SEM; n = 3). **C)** Dose response of the shift in voltage dependent activation (ΔV_0.5_) of Kv7.1/KCNE1 channels in the presence of DHA-taurine. **D)** Dose response of the change in maximal conductance (G_max_) of Kv7.1/KCNE1 channels in the presence of DHA-taurine. **E)** Current-voltage relationship of DHA-taurine on Cav1.2 channels (mean ± SEM; n = 3). **F)** Dose response of the shift in voltage dependent inactivation (ΔV_0.5_) of Cav1.2 channels in the presence of DHA-taurine. **G)** Dose response of the change in maximal conductance (G_max_) of Cav1.2 channels in the presence of DHA-taurine. **H)** Current-voltage relationship of DHA-taurine on Nav1.5 channels (mean ± SEM; n = 3). **I)** Dose response of the shift in voltage dependent inactivation (ΔV_0.5_) of Nav1.5 channels in the presence of DHA-taurine. **J)** Dose response of the change in maximal conductance (G_max_) of Nav1.5 channels in the presence of DHA-taurine.

These results suggest that PUFA analogues with taurine head groups exhibit broad selectivity for multiple ion channels.

### PUFA analogues with glycine head groups tend to be more selective for Kv7.1/KCNE1 with lower affinity for Cav1.2 and Nav1.5

PUFA analogues with glycine head groups have also been shown to effectively activate the Kv7.1/KCNE1 channel (37). The glycine head group has a lower pKa than regular PUFAs with a carboxyl head group (37), thereby allowing the head group to be more deprotonated and partially negatively charged at physiological pH. For this reason, PUFA analogues with glycine head groups are able to electrostatically activate Kv7.1/KCNE1 channels (37). We here tested several glycine compounds on Kv7.1/KCNE1, Cav1.2, and Nav1.5 channels to determine whether they have selective or non-selective effects on cardiac ion channels.

We first examined Linoleoyl glycine (Lin-glycine). Lin-glycine promotes the activation of the cardiac Kv7.1/KCNE1 channel by left-shifting the voltage dependence of channel activation to more negative voltages at 7 μM (−23.8 ± 1.6 mV; p < 0.001). Application of Lin-glycine also increases the G_max_ of Kv7.1/KCNE1 at 7 μM (2.3 ± 0.2; p = 0.008) (Fig. 8A-D). Lin-glycine inhibits Cav1.2 in a dose-dependent manner (Fig. 8E), but does not left shift the voltage dependence of inactivation (0.6 ± 2.3 mV; p = 0.65). Instead, Lin-glycine produces a decrease of the G_max_ at 7 μM, but this decrease is not statistically significant (0.3 ± 0.2; p = 0.07) (Fig. 8F-G). Lin-glycine causes a dose-dependent decrease of Nav1.5 current, left-shifts the voltage dependence of inactivation (−15.2 ± 2.8 mV; p = 0.01), and reduces the G_max_ at 7 μM (0.5 ± 0.1; p = 0.007) (Fig. 8H-J).

**Figure 8:**
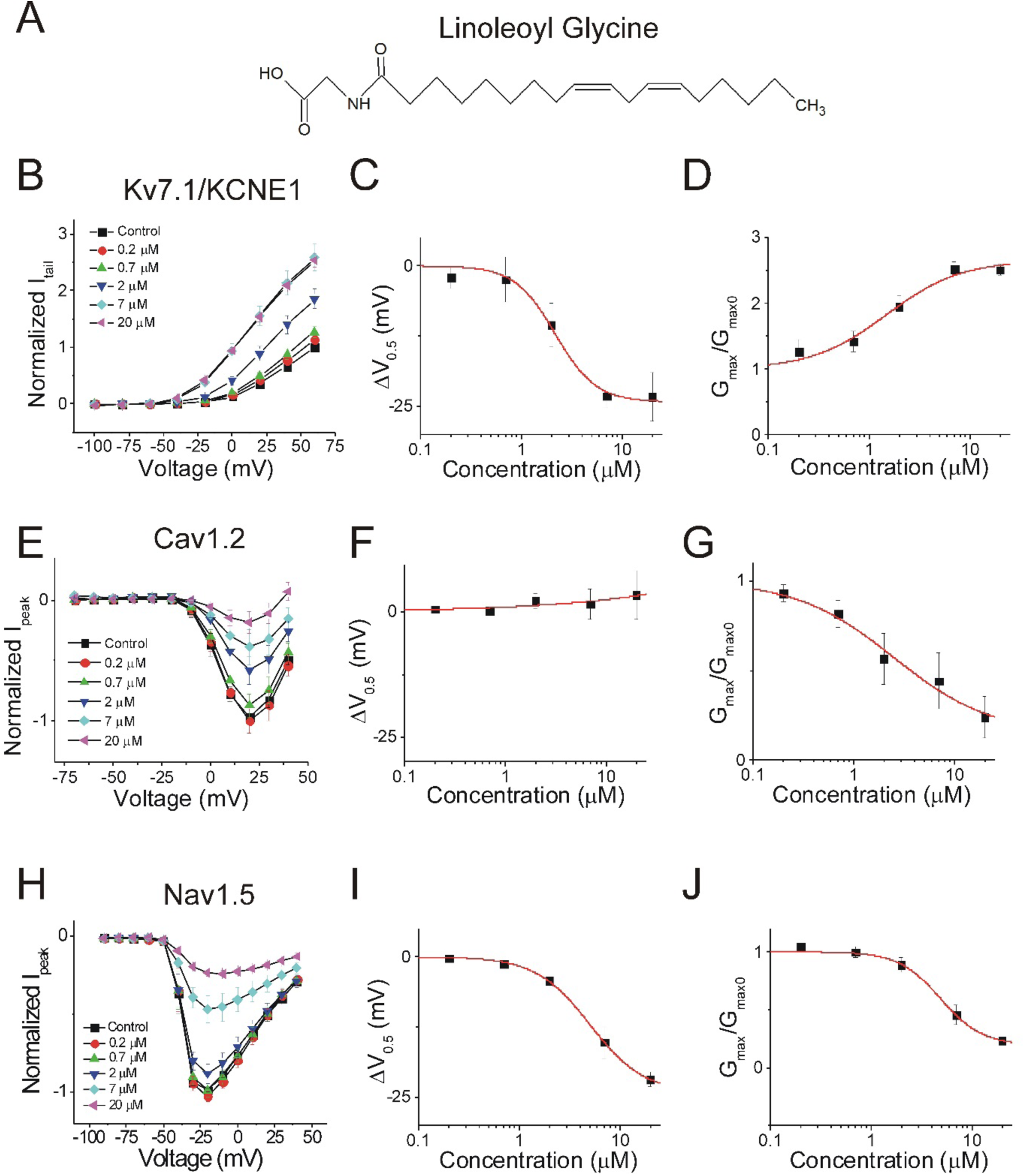
Linoleoyl glycine has broad selectivity for Kv7.1/KCNE1, Cav1.2, and Nav1.5. **A)** Structure of Linoleoyl glycine (Lin-glycine). **B)** Current-voltage relationship of lin-glycine on Kv7.1/KCNE1 channels (mean ± SEM; n = 4). **C)** Dose response of the shift in voltage dependent activation (ΔV_0.5_) of Kv7.1/KCNE1 channels in the presence of lin-glycine. **D)** Dose response of the change in maximal conductance (G_max_) of Kv7.1/KCNE1 channels in the presence of Lin-glycine. **E)** Current-voltage relationship of lin-glycine on Cav1.2 channels (mean ± SEM; n = 4). **F)** Dose response of the shift in voltage dependent inactivation (ΔV_0.5_) of Cav1.2 channels in the presence of lin-glycine. **G)** Dose response of the change in maximal conductance (G_max_) of Cav1.2 channels in the presence of lin-glycine. **H)** Current-voltage relationship of Lin-glycine on Nav1.5 channels (mean ± SEM; n = 4). **I)** Dose response of the shift in voltage dependent inactivation (ΔV_0.5_) of Nav1.5 channels in the presence of lin-glycine. **J)** Dose response of the change in maximal conductance (G_max_) of Nav1.5 channels in the presence of lin-glycine.

Pinoleoyl glycine (Pin-glycine) promotes the activation of Kv7.1/KCNE1 channels in a dose-dependent manner (Fig. 9A-B). Pin-glycine induces a left-shift in the voltage-dependence of activation (−8.7 ± 1.8 mV; p = 0.04) and increases the G_max_ at 7 μM (1.7 ± 0.1; p = 0.03) (Fig. 9C-D). Pin-glycine, however causes little inhibition of Cav1.2 current (Fig. 9E). Pin-glycine produces no shift in the voltage dependence of inactivation (−3.1 ± 4.2 mV; p = 0.54) and does not significantly reduce the G_max_ of Cav1.2 channels at 7 μM (0.8 ± 0.1; p = 0.34) (Fig. 9F-G). Pin-glycine inhibits Nav1.5 channels in a dose-dependent manner, but does not produce a significant left-shift in the voltage dependence of inactivation at 7 μM (−4.7 ± 1.9 mV; p = 0.09). However, there is a statistically significant reduction in the G_max_ at 7 μM (0.7 ± 0.08; p = 0.03) (Fig. 9I-J).

**Figure 9:**
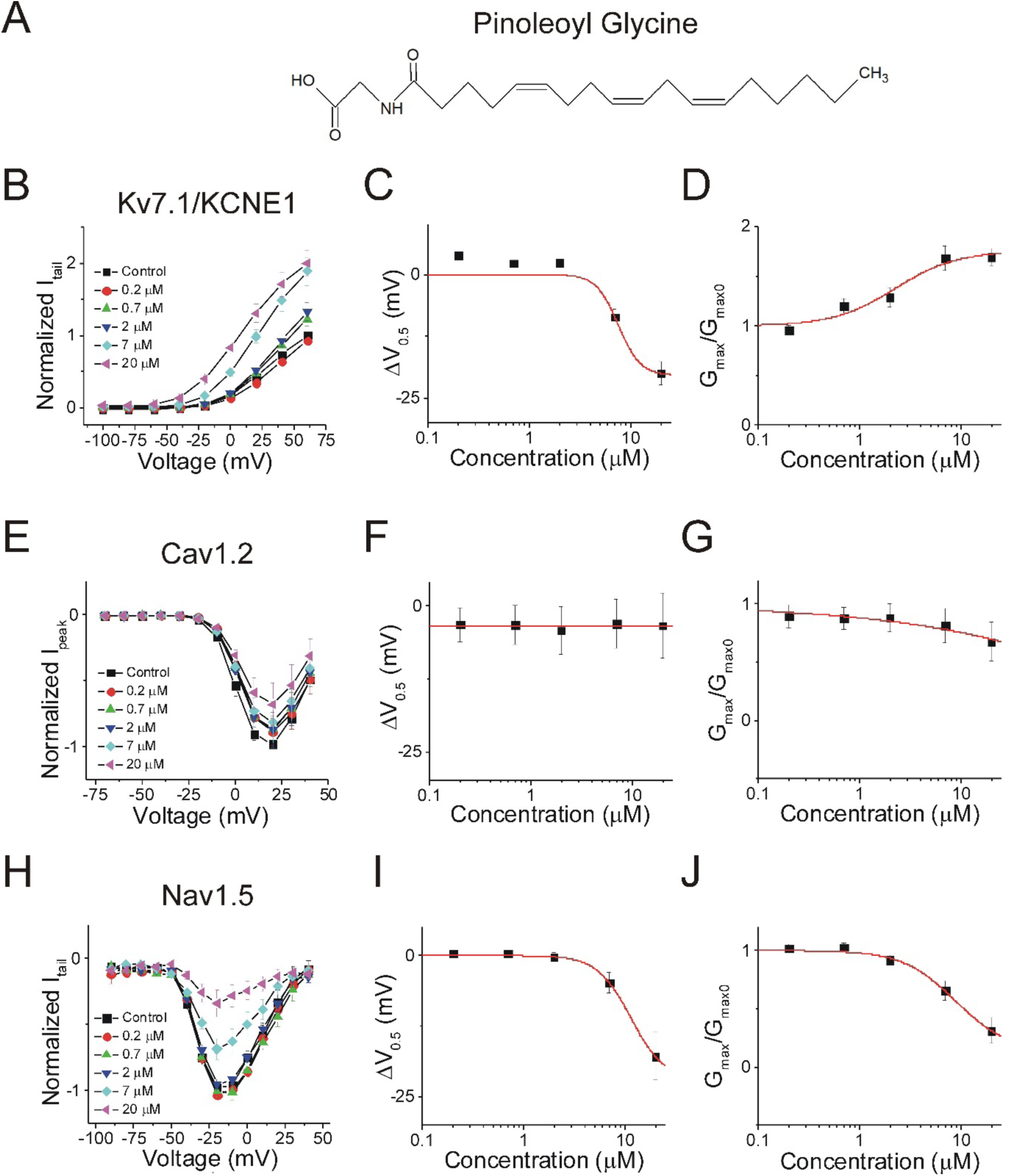
Pinoleoyl glycine is more selective for Kv7.1/KCNE1 and Nav1.5 channels than for Cav1.2. **A)** Structure of Pinoleoyl glycine (Pin-glycine). **B)** Current-voltage relationship of pin-glycine on Kv7.1/KCNE1 channels (mean ± SEM; n = 3). **C)** Dose response of the shift in voltage dependent activation (ΔV_0.5_) of Kv7.1/KCNE1 channels in the presence of pin-glycine. **D)** Dose response of the change in maximal conductance (G_max_) of Kv7.1/KCNE1 channels in the presence of Pin-glycine. **E)** Current-voltage relationship of pin-glycine on Cav1.2 channels (mean ± SEM; n = 3). **F)** Dose response of the shift in voltage dependent inactivation (ΔV_0.5_) of Cav1.2 channels in the presence of pin-glycine. **G)** Dose response of the change in maximal conductance (G_max_) of Cav1.2 channels in the presence of Pin-glycine. **H)** Current-voltage relationship of pin-glycine on Nav1.5 channels (mean ± SEM; n = 4). **I)** Dose response of the shift in voltage dependent inactivation (ΔV_0.5_) of Nav1.5 channels in the presence of pin-glycine. **J)** Dose response of the change in maximal conductance (G_max_) of Nav1.5 channels in the presence of pin-glycine.

DHA-glycine promotes the dose-dependent activation of Kv7.1/KCNE1 channels (Fig. 10A-B), left-shifting the voltage-dependence of activation (−10.5 ± 1.0 mV; p 0.002), and increasing the G_max_ at 7 μM (1.9 ± 0.2; p = 0.03) (Fig. 10C-D). However, DHA-glycine does not result in a dose-dependent decrease in Ca^2+^ currents (Fig. 10E). DHA-glycine does not left-shift the voltage dependence of inactivation (7.6 ± 3.1 mV; p = 0.13) and does not significantly decrease the G_max_ of Cav1.2 at 7 μM (1.0 ± 0.1; p = 0.98) (Fig. 10G). In addition, DHA-glycine produces some inhibition of Nav1.5, but only when applied at 20 μM (Fig. 10H). While DHA-glycine produces a small, but significant left-shift in voltage dependent inactivation at 7 μM (−4.8 ± 1.9 mV; p = 0.01), it does not significantly reduce the G_max_ at 7 μM (0.7 ± 0.08; p = 0.84) (Fig. 10H-J).

**Figure 10:**
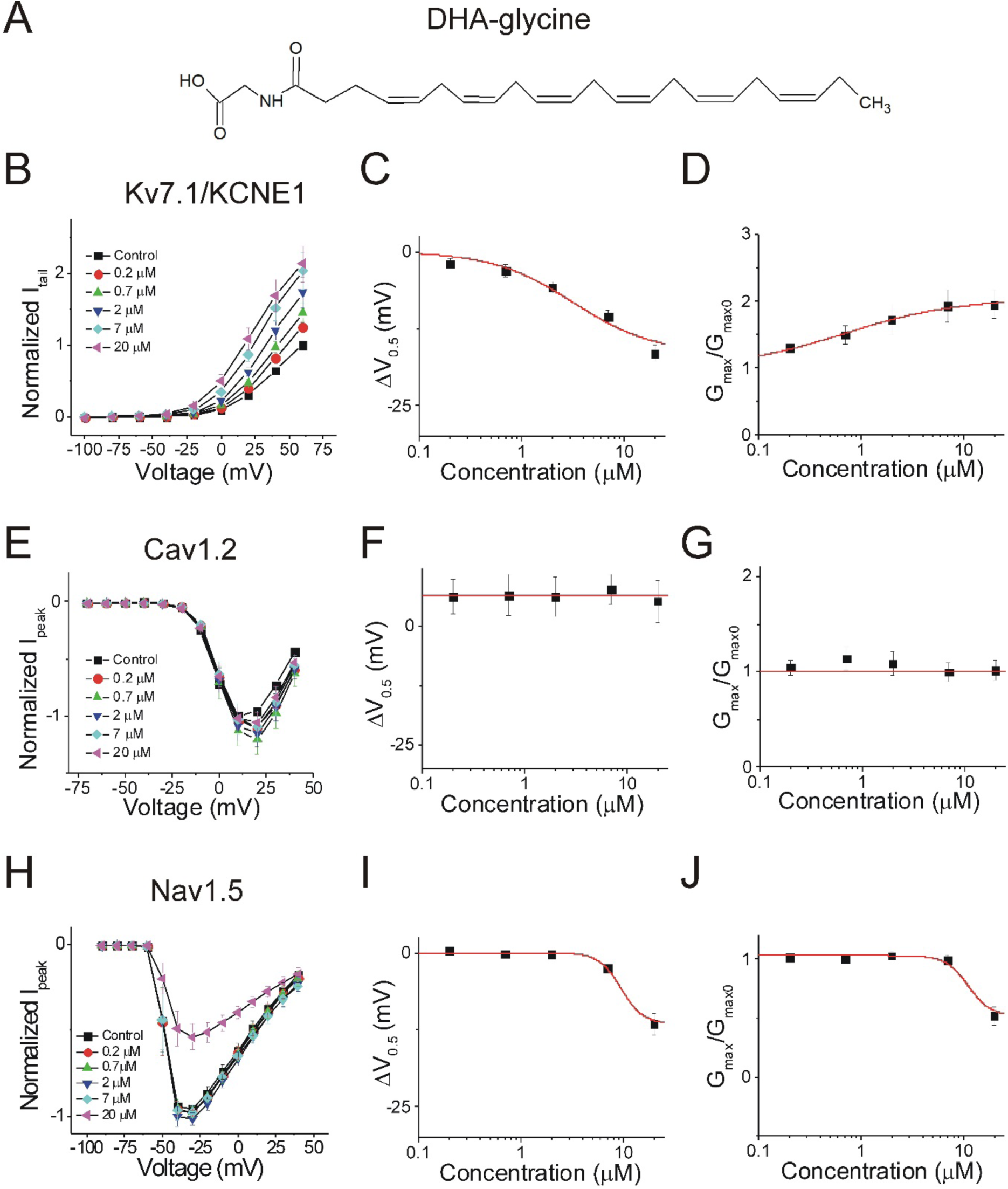
Docosahexanoyl glycine is more selective for Kv7.1/KCNE1 channels. **A)** Structure of docosahexanoyl glycine (DHA-glycine). **B)** Current-voltage relationship of DHA-glycine on Kv7.1/KCNE1 channels (mean ± SEM; n = 4). **C)** Dose response of the shift in voltage dependent activation (ΔV_0.5_) of Kv7.1/KCNE1 channels in the presence of DHA-glycine. **D)** Dose response of the change in maximal conductance (G_max_) of Kv7.1/KCNE1 channels in the presence of DHA-glycine. **E)** Current-voltage relationship of DHA-glycine on Cav1.2 channels (mean ± SEM; n = 3). **F)** Dose response of the shift in voltage dependent inactivation (ΔV_0.5_) of Cav1.2 channels in the presence of DHA-glycine. **G)** Dose response of the change in maximal conductance (G_max_) of Cav1.2 channels in the presence of DHA-glycine. **H)** Current-voltage relationship of DHA-glycine on Nav1.5 channels (mean ± SEM; n = 7). **I)** Dose response of the shift in voltage dependent inactivation (ΔV_0.5_) of Nav1.5 channels in the presence of DHA-glycine. **J)** Dose response of the change in maximal conductance (G_max_) of Nav1.5 channels in the presence of DHA-glycine.

These results suggest that PUFA analogues with glycine head groups tend to be more selective for the cardiac Kv7.1/KCNE1 channel and tend to have lower apparent affinity for Cav1.2 and Nav1.5 channels. Pin-glycine and DHA-glycine both have more selective effects on the Kv7.1/KCNE1 channel and lower apparent affinity for Cav1.2 and Nav1.5 channels compared to PUFA analogues with taurine head groups. Lin-glycine, however, is less selective for Kv7.1/KCNE1 channels compared to Pin-glycine and DHA-glycine. Lin-glycine modulates Kv7.1/KCNE1 and Cav1.2 at similar concentrations, while the modulatory effects of Lin-glycine on Nav1.5 take place at higher concentrations. This suggests that the combination of a glycine head group and linoleic acid tail boosts the apparent affinity for Cav1.2 and Nav1.5 channels.

### PUFA analogues with glycine head groups activate I_Ks_ with higher apparent affinity compared to I_Ca_ and I_Na_

We have observed that PUFA analogues have several different effects on the same channel (e.g. they alter voltage dependence and conductance at the same time). To evaluate the totally effects of PUFA analogues on channel currents at 0 mV (I/I_0_), we compared the dose response curves for I_Ks_ (Kv7.1/KCNE1), I_CaL_ (Cav1.2), and I_NaV_ (N_aV_1.5) (Table 1). At 0 mV, 7 μM N-AT does not increase I_Ks_ currents, but instead inhibits I_CaL_ currents and almost completely inhibits I_NaV_ currents (Fig. 11A; Table 1). By comparing the dose response curves and K_m_ (a measure of apparent binding affinity) for each channel current, we find that N-AT has similar apparent affinity for I_Ks_, I_CaL_, and I_NaV_. At 7 μM, Lin-taurine increases I_Ks_ (though not significantly) while significantly inhibiting I_CaL_ and I_NaV_, and exhibits higher apparent affinity for I_CaL_ and I_NaV_ than for I_Ks_ (Fig. 11B; Table 1). Similar to Lin-taurine, Pin-taurine and DHA-taurine increase I_Ks_ but it also inhibit I_CaL_ and I_NaV_ with higher apparent affinity for I_CaL_ and I_NaV_ than for I_Ks_ (Fig. 11C-D; Table 1). At 7 μM, Lin-glycine increases I_Ks_, but also inhibits I_CaL_ and I_NaV_ with higher apparent affinity for I_CaL_ than for I_Ks_ and I_NaV_ (Fig. 11E; Table 1). At 7 μM, Pin-glycine increases I_Ks_, but does not significantly inhibit I_CaL_ or I_NaV_. When we compare the K_m_ from the dose response curves of each channel current, we find that Pin-glycine has higher apparent affinity for I_Ks_ compared to I_NaV_ (Fig. 11F; Table 1). Lastly, at 7 μM, DHA-glycine increases I_Ks_ with little effect on I_CaL_ and I_NaV_, exhibiting higher apparent affinity for I_Ks_ and I_CaL_ than for I_NaV_ (Fig. 11G; Table 1). Overall, when we compare the effects of different PUFA analogues on I_Ks_, I_NaV_, and I_CaL_, PUFA analogues with taurine head groups tend to have higher apparent affinity for I_NaV_ and I_CaL_, whereas PUFA analogues with glycine head groups tend to have higher apparent affinity for I_Ks_.

**Figure 11:**
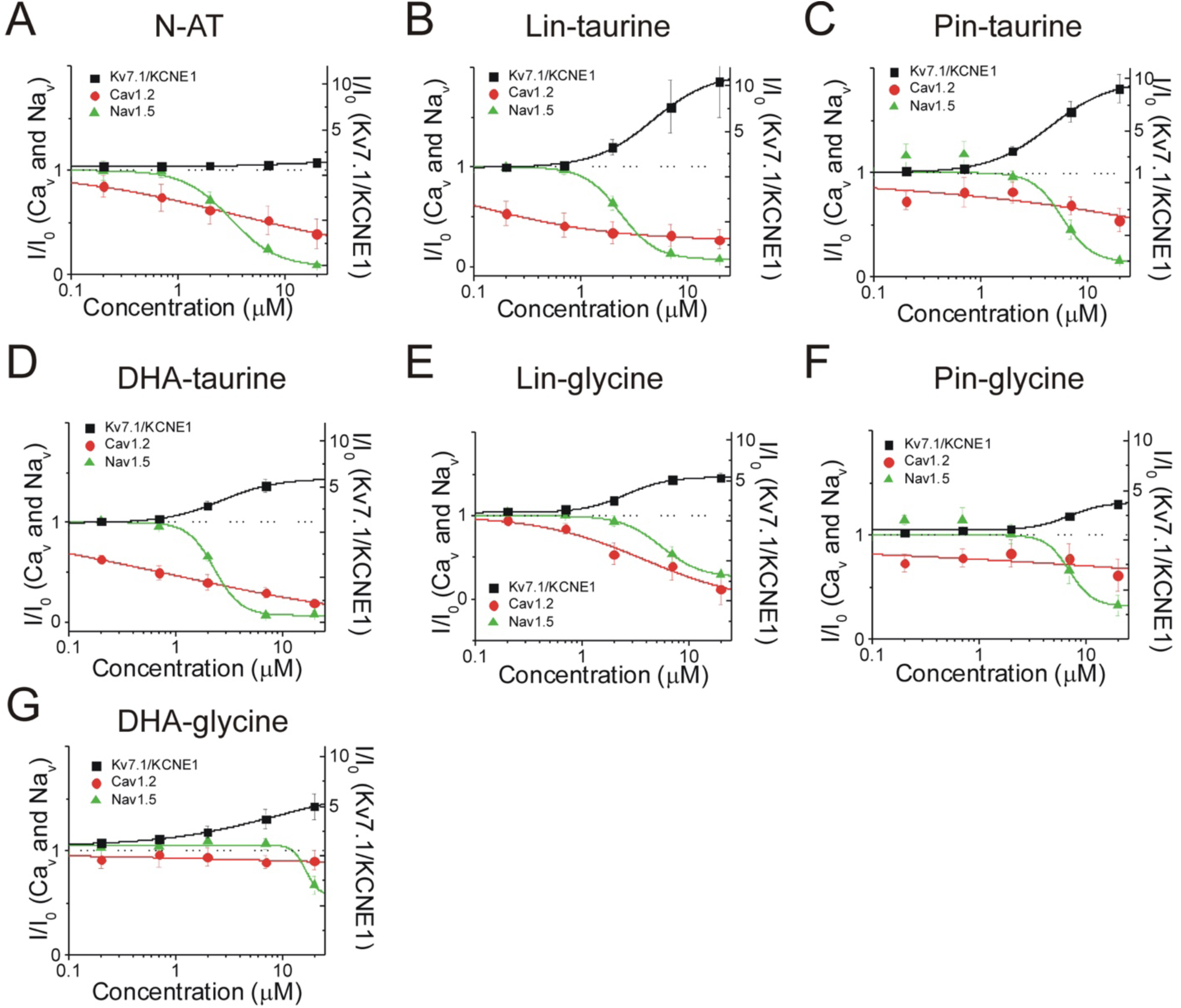
Dose response curves for PUFAs on I_Ks_, I_CaL_, and I_NaV_ at 0 mV. Dose response of **A)** N-AT, **B)** lin-taurine, **C)** pin-taurine, **D)** DHA-taurine, **E)** lin-glycine, **F)** pin-glycine, and **G)** DHA-glycine on I_Ks_, I_CaL_, and I_NaV_ currents (I/I_0_) at 0 mV.

**Table 1:**
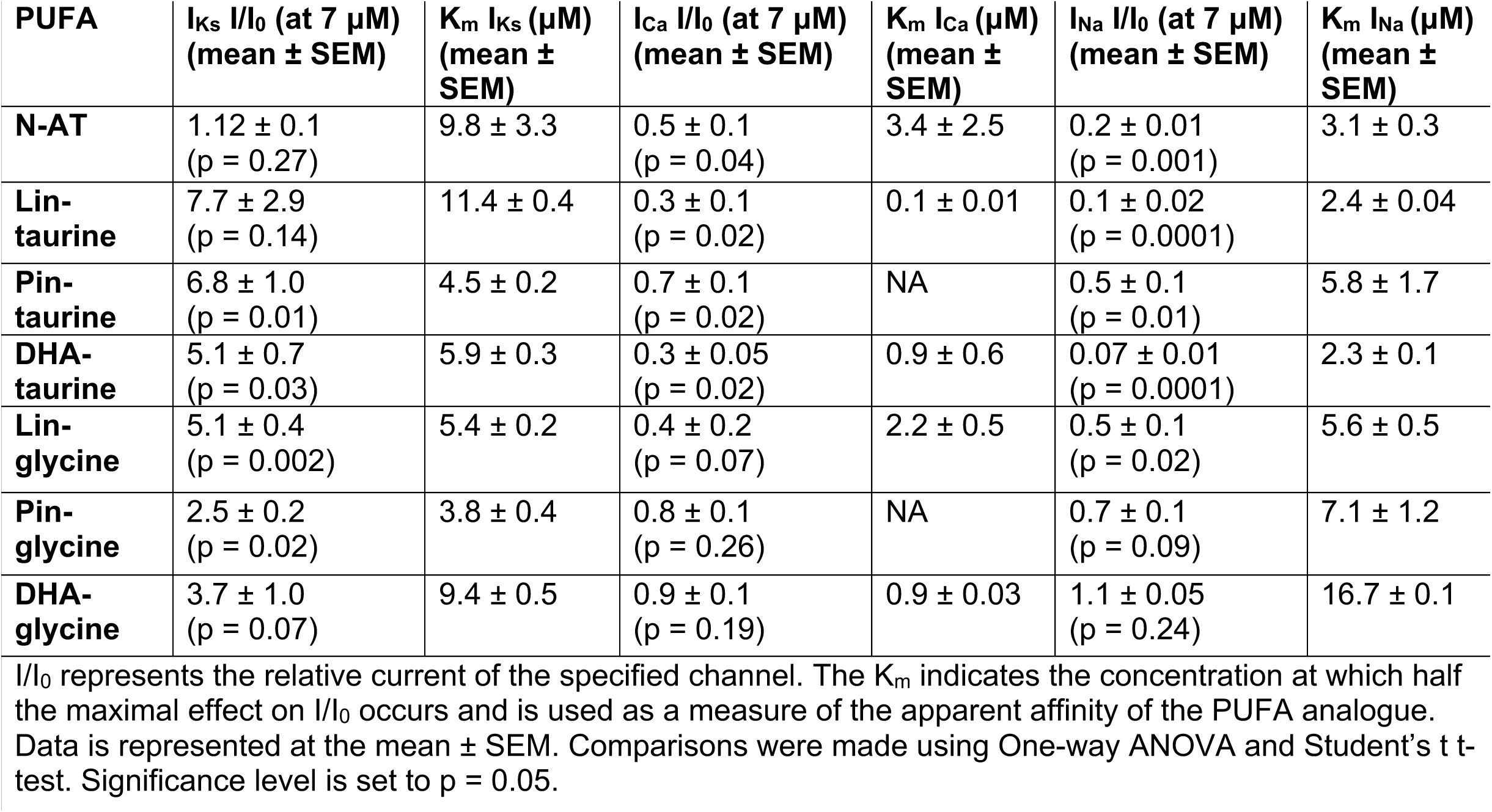
Summary of PUFA effects on current and apparent affinity

| PUFA | $I_{Ks} I/I_0$ (at 7 $\mu M$)<br>(mean ± SEM) | $K_m I_{Ks}$ ($\mu M$)<br>(mean ± SEM) | $I_{Ca} I/I_0$ (at 7 $\mu M$)<br>(mean ± SEM) | $K_m I_{Ca}$ ($\mu M$)<br>(mean ± SEM) | $I_{Na} I/I_0$ (at 7 $\mu M$)<br>(mean ± SEM) | $K_m I_{Na}$ ($\mu M$)<br>(mean ± SEM) |
| --- | --- | --- | --- | --- | --- | --- |
| **N-AT** | 1.12 ± 0.1<br>(p = 0.27) | 9.8 ± 3.3 | 0.5 ± 0.1<br>(p = 0.04) | 3.4 ± 2.5 | 0.2 ± 0.01<br>(p = 0.001) | 3.1 ± 0.3 |
| **Lin-**<br>**taurine** | 7.7 ± 2.9<br>(p = 0.14) | 11.4 ± 0.4 | 0.3 ± 0.1<br>(p = 0.02) | 0.1 ± 0.01 | 0.1 ± 0.02<br>(p = 0.0001) | 2.4 ± 0.04 |
| **Pin-**<br>**taurine** | 6.8 ± 1.0<br>(p = 0.01) | 4.5 ± 0.2 | 0.7 ± 0.1<br>(p = 0.02) | NA | 0.5 ± 0.1<br>(p = 0.01) | 5.8 ± 1.7 |
| **DHA-**<br>**taurine** | 5.1 ± 0.7<br>(p = 0.03) | 5.9 ± 0.3 | 0.3 ± 0.05<br>(p = 0.02) | 0.9 ± 0.6 | 0.07 ± 0.01<br>(p = 0.0001) | 2.3 ± 0.1 |
| **Lin-**<br>**glycine** | 5.1 ± 0.4<br>(p = 0.002) | 5.4 ± 0.2 | 0.4 ± 0.2<br>(p = 0.07) | 2.2 ± 0.5 | 0.5 ± 0.1<br>(p = 0.02) | 5.6 ± 0.5 |
| **Pin-**<br>**glycine** | 2.5 ± 0.2<br>(p = 0.02) | 3.8 ± 0.4 | 0.8 ± 0.1<br>(p = 0.26) | NA | 0.7 ± 0.1<br>(p = 0.09) | 7.1 ± 1.2 |
| **DHA-**<br>**glycine** | 3.7 ± 1.0<br>(p = 0.07) | 9.4 ± 0.5 | 0.9 ± 0.1<br>(p = 0.19) | 0.9 ± 0.03 | 1.1 ± 0.05<br>(p = 0.24) | 16.7 ± 0.1 |

$I/I_0$ represents the relative current of the specified channel. The $K_m$ indicates the concentration at which half the maximal effect on $I/I_0$ occurs and is used as a measure of the apparent affinity of the PUFA analogue. Data is represented at the mean ± SEM. Comparisons were made using One-way ANOVA and Student's t t-test. Significance level is set to p = 0.05.

### Selective Kv7.1/KCNE1 channel activators have antiarrhythmic effects in the simulated cardiomyocyte

We next tried to understand what kind of compound is the most effective at shortening the action potential duration. To determine whether selective PUFA analogues or non-selective PUFA analogues can shorten the action potential duration, we simulated the effects of applying the PUFA analogues on human cardiomyocyte using the O’Hara-Rudy dynamic (ORd) model (40) while modifying parameters for the voltage dependence and conductance for individual channels to reflect our experimental PUFA-induced effects. We simulated the effects of PUFA analogues that are non-selective modulators for cardiac ion channels (i.e. N-AT, Lin-taurine, Pin-taurine, DHA-taurine, and Lin-glycine) at concentrations of 0.7 μM, 2 μM, and 7 μM (Fig. 12A-E). In most cases, we saw little change in the ventricular action potential until we reached 7 μM where we were unable to elicit an action potential (Fig. 12A-E). One exception was the effect of applying DHA-taurine, in which case we observed a small shortening of the action potential at 0.7 μM, but then an abnormal action potential upstroke and prolongation of the action potential at 2 μM (Fig 12D). This is likely due to the potent block of Nav1.5 channels, causing the action potential to be largely calcium dependent. But again, at 7 μM DHA-taurine, we were unable to elicit an action potential (Fig. 12D). However, the PUFA analogues that were more selective for Kv7.1/KCNE1, such as Pin-glycine and DHA-glycine (at 7 μM) induce a slight shortening of the wild type ventricular action potential (Fig. 12F-G). For Pin-glycine and DHA-glycine, we induced Long QT Type 2 by simulating 25% block of the hERG channel, which generates the rapid component of the delated rectifier potassium current (I_Kr_). 25% hERG block prolongs the ventricular action potential by 50 ms. Application of Pin-glycine or DHA-glycine at 7 μM in the simulation partially restores the duration of the ventricular action potential. In addition to simulating the effects of PUFA analogues on the ventricular action potential duration, we also simulated the ability of 7 μM DHA-glycine (the most selective Kv7.1/KCNE1 activator) to prevent arrhythmia by simulating early afterdepolarizations using 0.1 μM dofetilide. Dofetilide is a blocker of the hERG channel and increases the susceptibility for early afterdepolarizations (40) (Fig. 12H). When we simulate 0.1 μM dofetilide + 7 μM DHA-glycine, we are able to suppress early afterdepolarizations, suggesting that the application of 7 μM DHA-glycine would be anti-arrhythmic (Fig. 12H).

**Figure 12:**
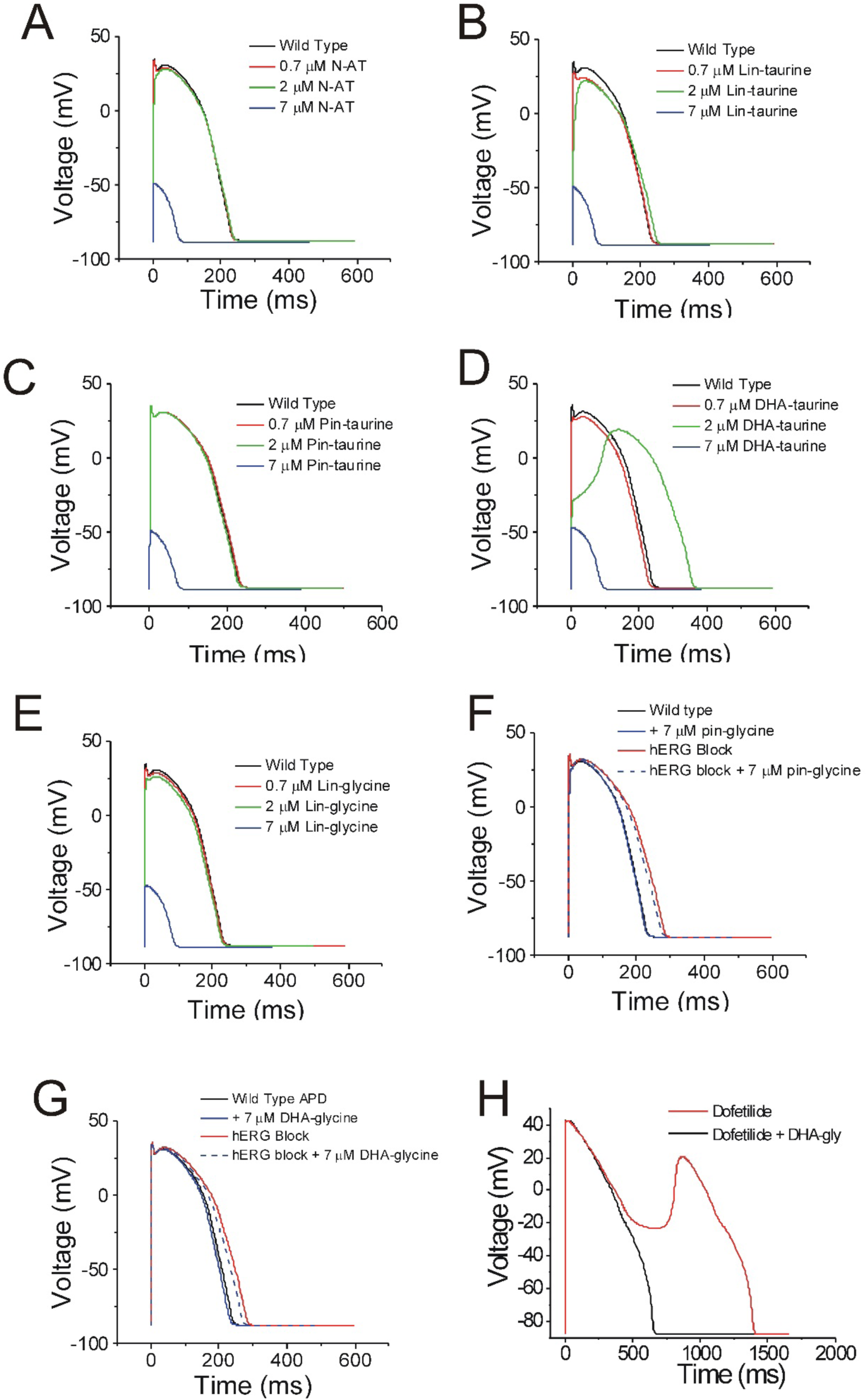
PUFAs that are selective for Kv7.1/KCNE1 channels partially restore prolonged ventricular action potential and suppress early afterdepolarizations. **A-G)** Simulated ventricular action potential in wild type cardiomyocytes (black) and in the presence of **A)** 0.7 (red), 2 (green), and 7 μM N-AT (blue), **B)** 0.7 (red), 2 (green), and 7 μM lin-taurine (blue), **C)** 0.7 (red), 2 (green), and 7 μM pin-taurine (blue), **D)** 0.7 (red), 2 (green), and 7 μM DHA-taurine (blue), **E)** 0.7 (red), 2 (green), and 7 μM lin-glycine (blue), **F)** 7 μM pin-glycine (blue solid), following 25% hERG block (red) and in the presence of 7 μM pin-glycine under 25% hERG block (blue dashed), and **G)** 7 μM DHA-glycine (blue solid), following 25% hERG block (red) and in the presence of 7 μM DHA-glycine under 25% hERG block (blue dashed). **H)** Early afterdepolarizations induced by dofetilide application (red) and suppression of early afterdepolarizations by 7 μM DHA-glycine in the presence of dofetilide (black).

## Discussion

We show here that PUFAs have different mechanisms of action on Kv7.1/KCNE1, Cav1.2, and Nav1.5 channels. We have previously shown that PUFAs promote the activation of Kv7.1/KCNE1 channels through the lipoelectric mechanism where the negatively charged PUFA head group electrostatically attracts both the S4 voltage sensor (facilitating its upward movement and channel opening) and K326 in S6 (increasing the maximal conductance) (36, 39, 42). In both Cav1.2 and Nav1.5 channels, PUFAs inhibit channel currents. We have found that PUFAs cause a dose-dependent reduction in the currents through in Cav1.2 channels, surprisingly with no effect on the voltage dependence of either activation or inactivation. In Nav1.5 channels, PUFAs cause inhibition through a dose-dependent decrease in currents, with both a left-shifting effect on the voltage dependence of inactivation and a decrease in conductance. We also demonstrate that PUFA analogues vary in their selectivity for voltage-gated ion channels. The selectivity depends on the specific concentration of PUFA applied, because several compounds have non-overlapping dose response curves for their effects on the three different channels (Fig. 11). We also found that PUFAs with taurine head groups tend to have broad modulatory effects on Kv7.1/KCNE1, Nav1.5 and Cav1.2 channels, with higher apparent affinity for Cav1.2 and Nav1.5 channels. Conversely, PUFAs with glycine head groups tend to be more selective for Kv7.1/KCNE1 channels and display lower apparent affinity for Cav1.2 and Nav1.5 channels. By understanding the effects of PUFA analogues on individual channels, it opens up the possibility to target specific forms of LQTS in which specific channels are mutated.

In this work, we have demonstrated that PUFA analogues modulate several different voltage-gated ion channels, including those underlying the ventricular action potential: Kv7.1/KCNE1, Nav1.5, and Cav1.2. The effects of PUFAs on Kv7.1/KCNE1, Cav1.2, and Nav1.5 individually are anticipated to have anti-arrhythmic effects and would potentially be beneficial for patients with Long QT Syndrome. In the case of I_Ks_ (Kv7.1/KCNE1) currents, PUFAs would be anti-arrhythmic by rescuing loss-of-function mutants of Kv7.1/KCNE1 (I_Ks_) channels in Long QT Type 1 (KCNQ1 mutations) or 5 (KCNE1 mutations). In the case of Na_v_ and Ca_v_ currents, PUFAs would be anti-arrhythmic by inhibiting gain-of-function mutants of Nav1.5 and Cav1.2 channels in Long QT Type 3 or 7, respectively. We used the O’Hara-Rudy Dynamic model to simulate the ventricular action potential in the presence of different PUFAs. In our simulations, PUFAs that are non-selective (i.e. that activate Kv7.1/KCNE1 while inhibiting Cav1.2 and Nav1.5) prevent the generation of an action potential. However, when we simulate the effects of Pin-glycine and DHA-glycine, which are both more selective for Kv7.1/KCNE1, we see a shortening in the action potential duration and the suppression of early afterdepolarizations. This suggests that selectively boosting I_Ks_ (Kv7.1/KCNE1) current would be important for shortening and terminating the ventricular action potential. However, evaluating PUFA-induced effects on Kv7.1/KCNE1, Cav1.2, and Nav1.5 channels bearing LQTS-causing mutations would be the next step in understanding the therapeutic potential for PUFA analogues as treatments for different forms of LQTS.

In our experiments using PUFA analogues on Nav1.5, we observed both a shift in the voltage dependence of inactivation and a dose-dependent decrease in Na^+^ currents. Extensive work has been done to characterize how each of the different voltage-sensing domains in Nav channels contribute to voltage-dependent activation and inactivation, many implicating DIV S4 in fast inactivation (6, 43). Recent work by Hsu and colleagues (2017) has also shown using voltage clamp fluorometry, the importance of both DIII and DIV in Nav channel inactivation (44). Our data suggest that PUFAs may interact with S4 segments involved in voltage-dependent inactivation, allowing PUFA analogues to left-shift the voltage dependence of inactivation. However, this does not completely explain the additional dose-dependent decrease in Na^+^ currents we observe on top of the leftward shifted voltage dependence of inactivation. Recent work by Nguyen and colleagues (2018) has uncovered a mechanism of Na_v_ channel inhibition through a new pathway, allowing a hydrophobic molecules to permeate a fenestration between domains III and IV (DIII and DIV) in the human cardiac Nav1.5 channel (45). It is possible that the hydrophobic PUFA analogue also block Nav1.5 channels through this fenestration between DIII and DIV, causing the voltage-independent decrease in sodium currents.

The molecular mechanism of action of PUFA analogues on Cav1.2 is still unclear, though we have shown that it does not occur through a shift in the voltage dependence of inactivation. In each case of Cav1.2 inhibition by PUFA analogues, we observe a dose-dependent decrease in the Ca^2+^ currents that appears as a linear decrease in I/I_0_ and G_max_. There is evidence that some Ca_v_ channel antagonists, such as dihydropyridines (DHPs) inhibit Ca_v_ channels through an allosteric mechanism (46). Pepe and colleagues (1994) found that DHA alters the effectiveness of dihydropyridines, suggesting a shared binding site, or nearby binding sites, for DHPs and PUFAs (47). Tang and colleagues (2016) found that dihydropyridines bind in a hydrophobic pocket near the pore of the bacterial Ca_v_Ab channel and cause an allosteric conformational change that leads to disruption of the selectivity filter and thus inhibition of Ca^2+^ currents (46). In addition, they observed that in the absence of DHPs a phospholipid occupies the DHP binding site (46). This would suggest that it is possible that PUFA analogues inhibit Cav1.2 by binding to, or near, the DHP binding site and causing an allosteric conformational change that leads to a collapse of the pore and thus explaining the inhibition of Cav1.2 currents without any changes in the voltage dependence of inactivation.

Work from several groups has demonstrated a shared electrostatic mechanism of action on voltage-gated K^+^ channels (48) and voltage-gated Na^+^ channels (49) by biaryl sulfonamides. Liin and colleagues (2018) showed that biaryl sulfonamides promote the activation of the Shaker K^+^ channel through an electrostatic effect on the voltage sensing domain. In addition, Ahuja and colleagues (2015) showed that aryl sulfonamides inhibit Na_v_ channels through an electrostatic “voltage sensor trapping” mechanism that is specific for the Nav1.7 isoform. The work by Ahuja et al. supports the ability to pharmacologically target different ion channels with a high degree of selectivity (49). This is in agreement with our findings using PUFA analogues that show that PUFA analogues are variable in their channel selectivity, allowing us to target particular ion channels involved in the ventricular action potential.

Our experiments were conducted using the *Xenopus laevis* oocyte expression system, where voltage-clamp recordings were performed at room temperature. It is possible that there may be temperature differences in the ways PUFA analogues modify different ion channels that we are unable to capture by conducting experiments at 20°C. There is also the possibility that the membrane composition may differ between *Xenopus* oocytes and mammalian cells or cardiomyocytes. However, using *Xenopus* oocytes, we are able to measure distinct differences between mechanisms of PUFA modulation in Kv7.1/KCNE1, Cav1.2, and Nav1.5 in isolation. To further confirm our findings, experiments should be conducted in mammalian cells or cardiomyocytes to determine the effects of different PUFA analogues on individual ion channels and the duration of the ventricular action potential at physiological temperatures.

The work presented here demonstrates that PUFA analogues exert diverse modulatory effects on different types of voltage-gated ion channels through non-identical mechanisms. Because PUFA analogues modulate Kv7.1/KCNE1 channels through electrostatic effects, we hypothesized they would have similar effects on Cav1.2 and Nav1.5 channels. However, our data suggests that PUFA analogues can exert various modulatory effects on the activity of different ion channels, and that the mechanism depends on the ion channel that is being modulated. In addition, we have shown that PUFA analogues exhibit a range of selectivity for different ion channels, which depends both on the PUFA head group and the combination of PUFA head and tail groups. Using simulations of the ventricular action potential, we have shown that selective Kv7.1/KCNE1 channel activators are the most effective at shortening a prolonged ventricular action potential and suppressing early afterdepolarizations induced by hERG block. This suggests that boosting Kv7.1/KCNE1 currents by using selective Kv7.1/KCNE1 channel activators can aid in restoring a normal action potential duration and possess antiarrhythmic potential.

## Methods

### Molecular Biology

cRNA encoding Kv7.1 and KCNE1, Nav1.5 and β1, and Cav1.2, β3, and α2δ were transcribed using the mMessage mMachine T7 kit (Ambion). 50 ng of cRNA was injected into defolliculated *Xenopus laevis* oocytes (Ecocyte, Austin, TX): For Kv7.1/KCNE1 channel expression, we injected a 3:1, weight:weight (Kv7.1:KCNE1) cRNA ratio. For Nav1.5 channel expression, we injected a 2:1, weight:weight (N_aV_1.5:β1) cRNA ratio. For Cav1.2 channel expression, we injected a 2:1:1, weight:weight (Cav1.2:β3:α2δ) cRNA ratio. Injected cells were incubated for 72-96 hours in standard ND96 solution (96 mM NaCl, 2 mM KCl, 1 mM MgCl_2_, 1.8 mM CaCl_2_, 5 mM HEPES; pH = 7.5) containing 1 mM pyruvate at 16°C prior to electrophysiological recordings.

### Two-electrode voltage clamp (TEVC)

*Xenopus laevis* oocytes were recorded in the two-electrode voltage clamp (TEVC) configuration. Recording pipettes were filled with 3 M KCl. The recording chamber was filled with ND96 (96 mM NaCl, 2 mM KCl, 1 mM MgCl_2_, 1.8 mM CaCl_2_, 5 mM HEPES; pH 7.5). For Cav1.2 channel recordings, *Xenopus* oocytes were injected with 50 nl of 100 mM EGTA and incubated at 10°C for 30 minutes prior to electrophysiological recordings in order to sequester cytosolic calcium. In addition, Cav1.2 channel recordings were done in Ca^2+^-free solutions, using Ba^2+^ as the charge carrier, to prevent calcium-dependent inactivation of Cav1.2 channels. PUFAs were obtained from Cayman Chemical (Ann Arbor, MI.) or synthesized in house (Linköping, Sweden) through methods previously described (Bohannon et al., submitted) and kept at −20°C as 100 mM stock solutions in ethanol. Serial dilutions of the different PUFAs were prepared from stocks to make 0.2 μM, 0.7 μM, 2 μM, 7 μM, and 20 μM concentrations in ND96 solutions (pH = 7.5). PUFAs were perfused into the recording chamber using the Rainin Dynamax Peristaltic Pump (Model RP-1) (Rainin Instrument Co., Oakland, CA. USA).

Electrophysiological recordings were obtained using Clampex 10.3 software (Axon, pClamp, Molecular Devices). To measure Kv7.1/KCNE1 currents we apply PUFAs as the membrane potential is stepped every 30 sec from −80 mV to 0 mV for 5 seconds before stepping to −40 mV and back to −80 mV to ensure that the PUFA effects on the current at 0 mV reached steady state. A voltage-step protocol was used to measure the current vs. voltage (I-V) relationship before PUFA application and after the PUFA effects had reached steady state for each concentration of PUFA. Cells were held at −80 mV followed by a hyperpolarizing prepulse to −140 mV. The voltage was then stepped from −100 to 60 mV (in 20 mV steps) followed by a subsequent voltage step to −20 mV to measure tail currents before returning to the −80 mV holding potential. For Cav1.2 channel recordings, PUFAs are applied as the membrane potential is stepped from −80 mV to −30 mV and then 10 mV before returning to the holding potential of −80 mV. This allows the PUFA effects to reach steady state before recording voltage-dependent activation and inactivation. To measure voltage-dependent activation of Cav1.2, cells are held again at −80 mV and then stepped from −70 mV to 40 mV (in 10 mV steps). Voltage-dependent inactivation was measured by holding cells at −80 mV, applying a 500-ms conditioning prepulse at voltages between −80 mV and 20 mV (in 10 mV steps) before stepping to a test pulse of 10 mV to measure the remaining current and returning to −80 mV holding potential. For Nav1.5 channel recordings, PUFAs are applied as the membrane potential is stepped from −80 mV to −90 mV for 480 ms before stepping to 30 mV for 50 ms and returned to a holding potential of −80 mV. This allows the PUFA effects to reach steady state before recording voltage-dependent activation and inactivation. To measure voltage-dependent activation of Nav1.5, cells are held at −80 mV and then stepped from −90 mV to 40 mV (in 10 mV steps) and then returning to −80 mV holding potential. Voltage-dependent inactivation was measured by holding cells at −80 mV, applying a 500-ms conditioning prepulse at voltages between −140 mV and −30 mV (in 10 mV steps) and measuring the remaining current at a test pulse of −30 mV before returning to −80 mV holding potential.

### Data analysis

Tail currents from Kv7.1/KCNE1 measures were analyzed using Clampfit 10.3 software in order to obtain conductance vs. voltage (G-V) curves. The V_0.5_, the voltage at which half the maximal current occurs, was obtained by fitting the G-V curves from each concentration of PUFA with a Boltzmann equation:

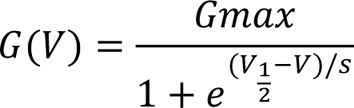

where G_max_ is the maximal conductance at positive voltages and s is the slope factor in mV. The current values for each concentration at 0 mV (I/I_0_) were used to plot the dose response curves for each PUFA. These dose response curves were fit using the Hill equation in order to obtain the K_m_ value for each PUFA:

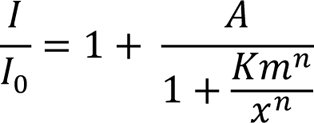

where A is the fold increase in the current caused by the PUFA at saturating concentrations, K_m_ is the apparent affinity of the PUFA, and n is the Hill coefficient. The maximum conductance (G_max_) was C_aL_culated by taking the difference between the maximum and minimum current values (using the G-V curve for each concentration) and then normalizing to control solution (0 μM). In Cav1.2 and Nav1.5 channels, peak currents (normalized to the peak values in control ND96) were used to determine PUFA induced changes in I/I_0_, ΔV_0.5_ of inactivation, and G_max_. Graphs plotting I/I_0_, ΔV_0.5_, G_max_, and K_m_ were generated using the Origin 9 software (Northampton, MA.). To determine if there were significant differences between apparent binding affinity of individual PUFA analogues for Kv7.1/KCNE1, Cav1.2, or Nav1.5 we conducted One-way ANOVA followed by Tukey’s HSD for multiple comparisons when comparing all three channels or Student’s t-test when comparing the apparent affinity for two channels. To determine if the PUFA-induced effects on I/I_0_, *Δ*V_0.5_, or G_max_ were statistically significant we conducted Student’s t-test on the PUFA-induced effects at 7 μM. Significance α-level was set at p < 0.05 – asterisks denote significance: p < 0.05*, p < 0.01**, p < 0.001***, p < 0.0001****.

### Simulations

The effects of individual PUFA analogues were simulated on each ion channel using Berkeley Madonna modeling software and equations from the MATLAB code in the O’Hara and Rudy Dynamic (ORd) model (40). We individually simulated the Kv7.1/KCNE1, Cav1.2, and Nav1.5 channels in Madonna and altered the parameters suggested to be modulated by PUFA binding to recapitulate our voltage clamp data from *Xenopus* oocytes. For example, to model the effects observed on the cardiac I_Ks_ channel, we modified the voltage dependence of channel activation by shifting the V_0.5_ as well as multiplying the I_Ks_ conductance by the factor increase we observed in our experiments at a given PUFA concentration.

MATLAB simulations of the ventricular action potential in the epicardium of the heart were performed using the ORd model (40). To simulate the effects of PUFAs, we introduced the same modified parameters in the MATLAB code as we used to model the PUFA effects on the ionic currents in Berkeley Madonna. We made simultaneous changes to Kv7.1/KCNE1, Cav1.2, and Nav1.5 for a given PUFA and specific PUFA concentration to model the effects of different PUFA analogues on the ventricular action potential under wild type and LQTS conditions. To simulate susceptibility to early afterdepolarizations, hERG block by 0.1 μM dofetilide was simulated which previously has been shown to cause spontaneous early afterdepolarizations (40). To simulate the ability of PUFA analogues to suppress early afterdepolarizations, we altered the activity of Kv7.1/KCNE1, Cav1.2, and Nav1.5 channels according to the PUFA-induced effects observed during experiments.

## Acknowledgements

We thank Levi Lindroos, Victor Kornfeld, Siri Lundholm, and Sankhero Gewarges for their contributions during their time as visiting scholars.

## Conflict of Interest

SIL, HPL: A patent application (62/032,739) has been submitted by the University of Miami with SIL and HPL as inventors. The other authors declare no other competing interests.

## Author Contributions

BMB, Acquisition of data, Analysis and interpretation of data, Drafting or revising the article; HPL, Conception and design, Analysis and interpretation of data, Drafting or revising the article; SIL, Conception and design, Acquisition of data, Analysis and interpretation of data, Drafting or revising the article; MEP, Acquisition of data; XW, Acquisition of data.

## Funding

This work was supported by NIH R01-HL131461 (to H. Peter Larsson) and by the Swedish Society for Medical Research and the Swedish Research Council (2017-02040) (to Sara I. Liin).

---

**Supplemental Figure 1:**
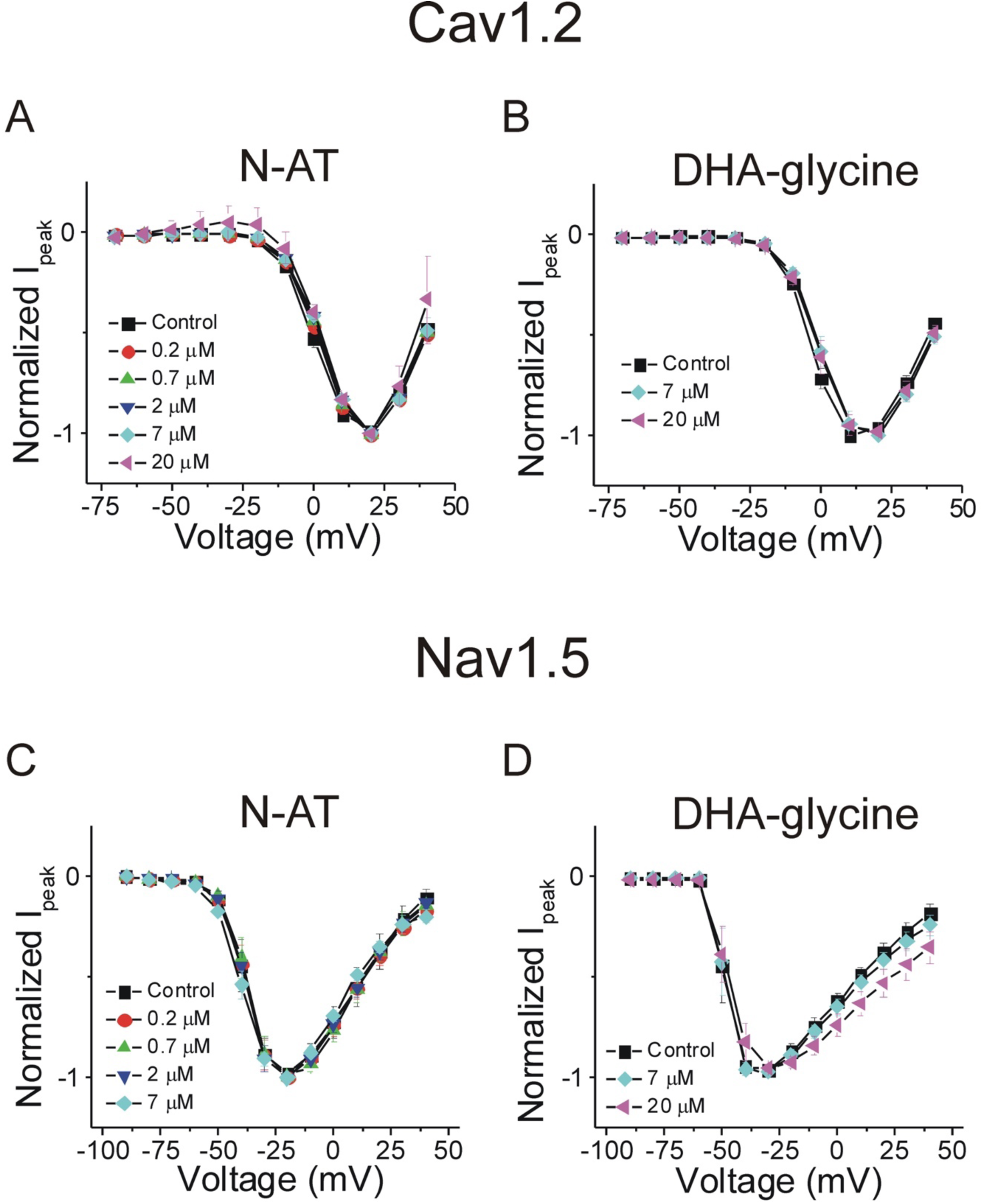
PUFA-induced changes in I/I_0_ normalized by concentration show no changes in voltage-dependent activation of Cav1.2 and Nav1.5 channels. **A-B)** Voltage-dependent activation of Cav1.2 in the presence of **A)** N-AT and **B)** DHA-glycine. Peak currents are normalized to each concentration to clearly visualize that there is no shifts in voltage-dependent activation. **C-D)** Voltage-dependent activation of Nav1.5 in the presence of **C)** N-AT and **D)** DHA-glycine. Peak currents are normalized to each concentration to clearly visualize that there are no shifts in voltage-dependent activation.

